# Determinants of missense pathogenicity in Usherin: Evaluating sequence and predicted-structure features

**DOI:** 10.64898/2026.04.23.720479

**Authors:** Diya Choudhary, Stephanie Portelli, David B. Ascher

## Abstract

USH2A encodes Usherin, a 5,202-residue multi-domain protein whose missense variation is a major cause of Usher syndrome type II. Interpretation remains challenging because the protein lacks a full-length experimental structure and thousands of reported missense variants remain unresolved. In an attempt to address this gap, we developed UshEffect-3D, a gene-specific machine-learning-based variant effect predictor that integrates sequence-based evolutionary measures, structure-based biochemical properties, local interaction terms, and predicted protein stability changes to prioritise USH2A missense variants of unknown significance (VUS). To ensure that the structure-derived features were extracted from reliably modelled predicted structures, the AlphaFold2 models of 47 annotated Usherin domains were compared against experimental structures of representative domain types. The predicted structures showed strong fold concordance with representative experimental structures, with 43 domains achieving TM-scores of at least 0.5. Among binary classifiers trained using a residue position-grouped cross-validation strategy, Logistic Regression was selected as the most stable performer and achieved an MCC of 0.75, AUC of 0.96, sensitivity of 0.93 and specificity of 0.81 on a group-aware held-out test set.

On a benchmarking task, UshEffect-3D performed comparably to six general-purpose variant-effect predictors evaluated on the held out test set, scoring the highest on every balanced metric but no statistically resolvable advantage over any major competitors such as PolyPhen-2, VESPAl, ESM-1b and AlphaMissense. Feature ablation displayed that sequence-derived evolutionary constraints were the sole detectable source of discrimination - withholding the conservation features reduced cross-validated MCC by 0.35 (paired 95% CI *−*0.45 to *−*0.25), whereas removing structural descriptors and predicted stability change negligibly affected model performance. Applying our model to the 2,639 ClinVar VUS, we reclassified 981 (37.2%) as likely pathogenic highlighting the extent of disease-relevant missense variation yet to be uncovered. Thus, UshEffect-3D serves as an interpretable protein-specific framework and a supplementary ranked resource that can be used to flag unresolved variants for further clinical and experimental review.

## Introduction

Interpreting the functional consequences of rare disease variants remains a central problem^1^, and it is especially acute for large, multi-domain proteins where full-length experimental structures are unavailable and functional understanding is sparse^2^. The rapid growth of clinical sequencing has outpaced the ability to resolve how individual amino acid substitutions perturb folding, local packing, and inter-domain or intermolecular interactions ^3,4^. Ideally, bridging this gap requires placing each substitution in its residue-level structural environment to exploit information that sequence-only predictors might miss - an approach that has previously worked well for some clinically important proteins such as p53^5^ and PTEN^6^.

*USH2A* encodes Usherin, a large, multi-domain protein. The canonical long isoform (UniProtKB: O75445^7,8^) comprises 5202 amino acids and more than 47 domains spanning Laminin, Laminin EGF-like, Laminin G-like, and Fibronectin type III repeats, in addition to a short transmembrane region and a PDZ-binding motif^7,9^ (Figure 1B). Usherin assembles into the USH2 complex with Adhesion G protein-coupled Receptor V1 (ADGRV1) and Whirlin, supporting photoreceptor maintenance and hair-bundle organization^10,11^. No full-length experimental structure exists^7,12^, so the structural basis of missense pathogenicity in this protein has been essentially inaccessible. This matters clinically because *USH2A* is among the most frequently mutated genes in inherited retinal disease and the leading cause of Usher syndrome type II (OMIM: 276901^13^), the most prevalent form of Usher syndrome^14^ and a common cause of combined hearing and vision loss^15–18^. Over 70% (*n* = 2639) of its ClinVar variants remain of uncertain significance (as of April 2026) ^19,20^. Usherin therefore combines large size, extensive domain repetition, and a high burden of uninterpreted variants - conditions under which one might expect structural context to be informative.

**Figure 1:**
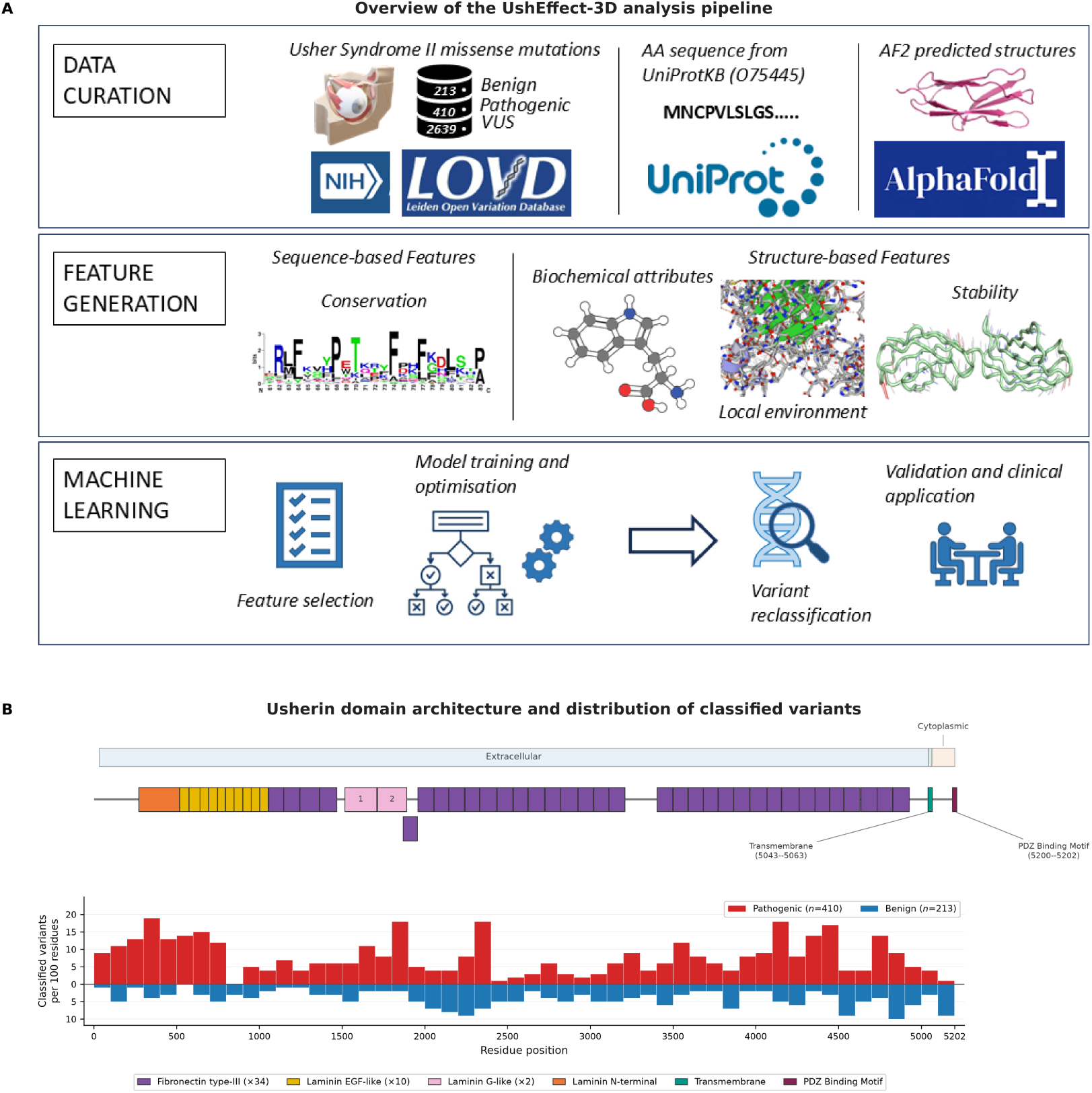
Overview of the UshEffect-3D methodology and the domain architecture of Usherin. **(A)** Schematic of the three-stage computational pipeline. *Data curation*: labelled missense variants (213 benign, 410 pathogenic) and 2639 variants of uncertain significance were retrieved from ClinVar and the Leiden Open Variation Database (LOVD), the canonical Usherin amino acid sequence (UniProtKB: O75445) was obtained from UniProtKB, and domain structures were predicted using AlphaFold2. *Feature generation*: 153 sequence- and structure-based features were computed per variant, spanning evolutionary conservation, biochemical attributes, local residue environment, and predicted thermodynamic stability. *Machine learning* : features were passed through an all-relevant selection pipeline (Spearman and variance-inflation filtering followed by Boruta), reducing the set to eleven descriptors, and eleven classifiers were then compared by nested cross-validation in which variants were grouped by residue position so that no position contributed to both training and evaluation. The selected model was the one with the highest mean nested cross-validation MCC after penalising fold-to-fold variability, and was evaluated on a group-aware held-out test set and on an independent set of 78 ACMG-classified pathogenic variants withheld from all development steps before being applied to the unresolved VUS. **(B)** Domain architecture of Usherin according to Uniprot, across the full 5202-residue sequence, with the distribution of classified variants below it. The upper lane marks membrane topology (extracellular, transmembrane and cytoplasmic spans). Blocks on the backbone are the 49 annotated domain instances, coloured by type: Fibronectin type III (purple, *×*34), Laminin EGF-like (yellow, *×*10), Laminin G-like (pink, *×*2), Laminin N-terminal (orange), transmembrane (teal) and PDZ binding motif (wine); the same colours are used for these families throughout. The transmembrane and PDZ blocks are drawn at a minimum visible width and are therefore not to scale. Fibronectin type III instance 5 (residues 1869–1955) is offset onto a second row because it overlaps Laminin G-like instance 2 by 23 residues. The lower track shows the 623 classified variants in fixed 100-residue windows, with pathogenic counts (red) above the baseline and benign counts (blue) below. Because the windows are of equal width, counts are directly comparable along the sequence. They are absolute counts rather than proportions.

Advances in protein structure prediction, notably AlphaFold2^21^, now recover high-confidence structures for monomers that remain experimentally intractable at full length^22^. This makes it possible to characterize each substitution by its predicted local environment, stability effect, and packing interactions^5,6,23^, and to test whether such descriptors capture the determinants of pathogenicity in a protein of this scale.

Here we present UshEffect-3D, a machine learning-based variant effect prediction framework which integrates AlphaFold2-based domain modeling^21^, sequence conservation^24^, and protein structural descriptors^25,26^ to map the pathogenicity landscape of *USH2A* missense variants (Figure 1A). We show that pathogenic variants preferentially occupy evolutionarily constrained positions, and that a gene-specific model built on the resulting descriptors performs comparably to leading general-purpose predictors while supplying a decision threshold calibrated to this gene. Although the AlphaFold2-derived domains recover their expected folds at high confidence, the structural descriptors computed from them add no detectable discrimination beyond sequence conservation for this protein. Thus, our approach broadly demonstrates how predicted structure might inform missense variant interpretation in large multi-domain proteins like Usherin, and highlights the need to test whether structural features add value beyond evolutionary context alone.

## Methods

### 0.1 Structural curation

Experimental structures of Usherin were unavailable^12^, so the structure was predicted with AlphaFold2 version 2.3^21^ using the canonical amino acid sequence (UniProtKB O75445^7,8^) as input. As the 5202-residue sequence exceeded AlphaFold2’s input-length limit (1200 residues at the time of analysis), it was divided into five overlapping fragments of 1200 residues, with 200-residue overlaps between consecutive fragments, and each fragment was predicted independently.

The predicted fragments were assembled into a single full-length model by multiple-template comparative modelling in MODELLER version [10.4]^27^, using all five fragments simultaneously as templates against the full-length target sequence (MODELLER’s `model_multiple.py` script in the `AutoModel` class). Under this scheme, residues covered by two fragments (the 200-residue overlaps) were modelled from spatial restraints contributed by both covering templates rather than that of a single fragment.

However, due to a lack of confidence in inter-domain positioning, we extracted individual domains, including unannotated regions, from the predicted structure based on annotations provided in the UniProtKB database^7,8^. The structures were extracted using PyMOL version 2.5.0^28^ and saved as separate PDB files for structural feature generation.

The presence of the PDZ binding site indicates that Usherin participates in the formation of protein-protein complexes^10,29^. However, we did not include complexes in our analysis because the structures of the proteins that bind to Usherin have not been experimentally characterised (at the time of writing) and are very large, posing similar challenges to the prediction of the Usherin structure. Moreover, not every target involved in the USH2 complex contains abundant variant information with reliable annotations. Therefore, for this study, we did not account for variants associated with ADGRV1 and Whirlin.

### 0.2 Assessment of domain fold concordance

Fold concordance of the AlphaFold2-derived Usherin domains^21^ was assessed by superimposing each predicted domain onto a representative experimental structure of the same fold family. High resolution templates were obtained from the Protein Data Bank Database^12^. Domain types were mapped to representative crystal structures as follows: Fibronectin type III to 2E3V (Fibronectin type III domain of human neural cell adhesion molecule from human muscle culture lambda-4.4; X-ray, resolution=1.95 Å)^30^, Laminin G-like to 3SH4 (Laminin G-like domain 3 of human Perlecan; X-ray, resolution=1.50 Å)^31^, and both Laminin EGF-like and Laminin N-terminal to 4OVE (mouse Netrin-1; X-ray, resolution=2.64 Å)^32^. Netrin-1 is a laminin-superfamily protein^32^ whose Laminin N-terminal (LN) domain and laminin-type EGF (LE) modules correspond to these two Usherin laminin domain families.

Each of the 47 domain instances was aligned to its assigned template with TM-align (version 20240303) ^33^. Structural agreement was quantified by the TM-score normalised to the length of the predicted domain (i.e. the query rather than the reference), because some of the templates - most notably the multi-domain laminin structure 4OVE (422 residues) - were substantially larger than the individual excised domains, rendering reference-length normalisation uninformative. A TM-score *≥* 0.5 was taken to indicate recovery of the correct fold, with *≥* 0.7 denoting close structural correspondence^34^. For each domain instance, the TM-score and the C*α* root-mean-square deviation (RMSD) over the structurally aligned residues were recorded directly from the TM-align output, then summarised per domain type as the median and interquartile range. An overall median TM-score was also computed across all 47 instances, weighted by the number of modelled residues per instance.

To visualise representative superpositions, the single instance whose TM-score was closest to the per-type median was selected for each domain type. For each representative, the TM-align rotation matrix (*−m* option) was applied to the predicted domain in PyMOL version 2.5.0^28^ so that the displayed superposition exactly reproduced the TM-align alignment used to derive the reported metrics.

#### 0.2.1 Per-residue model confidence

AlphaFold2 per-residue confidence (pLDDT) was extracted from each predicted chunk’s B-factor column in the PDB file. As consecutive fragments overlap by 200 residues, residues in overlaps have two independent pLDDT values. These were resolved by retaining the copy furthest from its own fragment terminus, on the basis that predictions degrade towards fragment edges where the sequence context is truncated^21^. Residues were assigned to domain families using domain ranges specified in Uniprot^7^, and pLDDT was summarised per domain family as the median, interquartile range, and the percentage of residues at pLDDT *≥* 70 and *≥* 90.

### 0.3 Variant curation

Missense variants associated with the *USH2A* (*n* = 623; accessed August 2025) gene were downloaded from the ClinVar database^19^ and the Leiden Open Variation Database (LOVD)^35^. Variants were labelled as “pathogenic” if they were classified as pathogenic/likely pathogenic (*n* = 410) and “benign” if they were classified as benign/likely benign (*n* = 213). All ClinVar mutations with conflicting interpretations of pathogenicity (*n* = 335) were excluded. An additional “VUS” list of unique variants labelled as variants of uncertain significance (*n* = 2639) was obtained from ClinVar^19^ (accessed April 2026). Duplicated instances of variants across both databases were removed if clinical classifications were consistent. A separate validation set of 78 pathogenic variants classified according to the American College of Medical Genetics and Genomics (ACMG) standards^36^ was shortlisted from LOVD^35^ and excluded from model training and hyperparameter tuning. After removing these validation variants, 545 labelled variants remained in the curated training dataset.

### 0.4 Distribution of variants across domain types

Variant density was computed as the number of variants observed in a domain type divided by the number of residues of that type, per 100 residues. Enrichment was assessed by comparing pathogenic against benign counts within each domain type to the remainder of the protein, summarised by an odds ratio with a 95% confidence interval and tested with Fisher’s exact test. The same comparison was applied to the VUS set using the model’s predicted classes. The PDZ binding motif and transmembrane segment were excluded from both analyses as they were too small to support inference. Thus, every odds ratio is conditioned on the set of variants excluding the 14 variants that lie in the transmembrane region and PDZ motif. (620 of the 623 classified variants and 2625 of the 2639 VUS). All *p* values were corrected across domain types by the Benjamini-Hochberg procedure, and enrichment is claimed only where the corrected *q* value falls below 0.05.

### 0.5 Data splitting

As multiple substitutions frequently occur at the same site, a conventional stratified split would allow different variants at one position to fall on both sides of the partition, leaking position-specific data into the test set. Therefore we split our training dataset into *train* (80%) and *test* (20%) using scikit-learn’s `StratifiedGroupKFold`^37^ strategy - grouping variants by residue position so that every position falls entirely within either the training or the test set while class balance is approximately preserved. This left us with 435 training variants and 110 test variants.

### 0.6 Feature generation

A total of *n* = 153 features were generated using sequence- and structure-based *in silico* biophysical tools to account for mutation effects on protein structure and function. Sequence features were calculated on the wild-type canonical amino acid sequence from UniProtKB (accession: O75445) ^7,8^ and structural features were computed on individual domain models.

1. Sequence-based Features:

- *Evolutionary conservation:* We calculated features from ConSurf^38^, which captures the rate of evolution of an amino acid position within a protein family; DeMaSk^24^, to compute the logarithm of the frequency of a mutant amino acid and Shannon entropy^39^; and MTR-Viewer^40^, which provides the missense tolerance ratio (MTR). Lastly, we obtained Position-Specific Independent Counts (PSIC)^41^ using Envision^42^.
- *General biochemical attributes:* Intrinsic disorder levels and disordered binding regions were determined using IUPred3^43^ and ANCHOR2^43^, respectively. We also included chemical properties pertaining to the wild-type and mutant amino acids, such as weight, volume, size, and isoelectric points^42^.
2. Structure-based Features:

- *Local interactions:* The wild-type residue environment was represented as a graph where mutated residues were nodes and distances were edges to generate graph-based signatures^25^. Arpeggio^26^ was used to calculate all intra- and interatomic interactions (e.g. van der Waals, hydrophobic, hydrogen-bond, ionic, aromatic, and covalent). These were calculated on both wild-type and mutant structures (generated via MODELLER 10.3^27^) to represent the environment before and after mutation. We also used CSM-potential^44^ to predict potential protein-protein interaction sites.
- *Stability:* To evaluate dynamic shifts, we utilised DDmut^45^, which captures the change in Gibbs Free Energy (ΔΔ*G*) following a mutation.

### 0.7 Feature selection

To ensure best practices, the feature selection pipeline was only applied to the training split, prior to cross validation. Pairwise Spearman correlation filtering was applied first to remove redundant features using the pandas `DataFramėcorr` function^46,47^, retaining the feature with the higher mutual information score with respect to the ClinVar Significance labels whenever two features exceeded a correlation threshold of 0.9. Variance inflation factor (VIF) filtering was then applied iteratively to the remaining features using the `variance_inflation_factor` function from the statsmodels package^48^, dropping the highest-VIF feature at each step until all remaining features had VIF *≤* 20. This was done to address multicollinearity that pairwise filtering alone cannot detect.

The python implementation of the Boruta algorithm^49^ was applied to the VIF-filtered feature set to identify all-relevant features with statistically significant importance relative to randomised shadow features. The selection relies on a balanced Random Forest base learner with parameters `max_depth=5` and `class_weight=’balanced’`. The `BorutaPy` algorithm was configured to `max_iter=50` and `n_estimators=’auto’`. This sequential feature selection pipeline reduced the original 153 features to a final set of eleven in the training set. The test set was also reduced to this final subset of features.

### 0.8 Normalisation

The features were standardised using the z-score normalisation method implemented using scikit-learn’s `StandardScaler` function^37^. It was fitted exclusively on the training set and subsequently applied to the test set.

### 0.9 Model training

Eleven classification algorithms were trained and evaluated. These include Extra Trees, Random Forest, Decision Tree, Gradient Boosting, XGBoost, AdaBoost, Gaussian Naive Bayes, K-Nearest Neighbours, Support Vector Classifier (SVC), Multilayer Perceptron (MLP), and Logistic Regression. The XGBoost model was based on the python-based XGBoost package^50^. All other models were implemented using scikit-learn^37^.

Model comparison used nested, position-grouped cross validation so that hyperparameter tuning could not contaminate the generalisation estimate. The outer loop was a 10-fold `StratifiedGroupKFold`^37^ over the training set. Within each outer training fold, hyperparameters were tuned independently by `RandomizedSearchCV` (`n_iter=30` configurations, scored by MCC) over an inner 5-fold `StratifiedGroupKFold` cross validation, and each resulting tuned estimator was scored on the outer fold test split.

Model selection was based on nested cross-validation results alone. Because the mean nested-CV MCC of the leading classifiers fell well within one standard deviation of one another, ranking on the mean alone would amount to selecting on noise. Models were therefore ranked by the mean nested-CV MCC minus its standard deviation across the ten outer folds, a criterion that rewards strong performance while penalising instability between folds. This selection rule was fixed before the test set was examined. Following selection, each classifier was retuned once on the complete training set through the same inner grouped cross-validation and refit with the best configuration (`refit=True`) for final evaluation. For the selected Logistic Regression classifier, the tuning returned an L2 penalty with *C* = 1.5 *×* 10*^−^*^3^ (`saga` solver, `class_weight=‘balanced’`, `max_iter=1000`), and therefore a strongly regularised fit.

### 0.10 Model evaluation

The scoring metric for tuning was the Matthews Correlation Coefficient (MCC). Per-fold performance metrics including MCC, accuracy, balanced accuracy, F1 score, precision, sensitivity, and specificity were computed using the `sklearn.metrics` module^37^.

MCC was computed as:

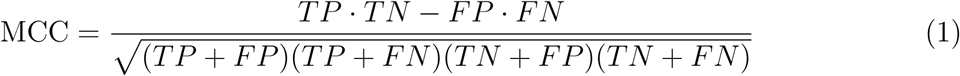

where *TP*, *TN*, *FP*, and *FN* denote the number of true positives, true negatives, false positives, and false negatives, respectively. MCC was selected as the primary evaluation metric over accuracy and F1 score due to the class imbalance present in the dataset, as it accounts for all four cells of the confusion matrix and returns a balanced score even when the classes are of substantially different sizes^51^. MCC ranges from *−*1 (total disagreement between prediction and observation) to +1 (perfect prediction), with a value of 0 corresponding to random classification^51^.

#### 0.10.1 Confusion matrices

Confusion matrices were computed for the top-3 classifiers on the blind test set using scikit-learn’s `confusion_matrix` function^37^. Matrices were row-normalised by true class to show the proportion of correctly and incorrectly classified variants within each class, with absolute counts also reported (Figure 3B). This allows direct comparison of false positive and true positive rates across models independent of class size.

#### 0.10.2 Threshold-independent performance curves

Receiver Operating Characteristic (ROC) and Precision-Recall (PR) curves were computed for the top three classifiers from the predicted probabilities (`predict_proba`) on the held-out test set, using the `roc_curve`, `auc`, `precision_recall_curve` and `average_precision_score` functions from scikit-learn^37^. Both summarise performance across all classification thresholds and are therefore independent of the 0.5 decision boundary scored by the confusion matrices and MCC. The area under the ROC curve (AUC) measures discriminative ability wherein the probability that a randomly chosen pathogenic variant is scored above a randomly chosen benign one, with 0.5 corresponding to a random classifier and 1.0 to perfect discrimination.

The PR curve describes the trade-off between precision and recall at different thresholds which helps us evaluate at what point recovering more pathogenic variants shifts towards a higher false positive rate. As precision is diluted by the number of negatives in the pool, its baseline for an uninformative classifier is the positive-class prevalence (0.609 in the held-out set) rather than 0.5. Both curves were reported because AUC is insensitive to class balance whereas average precision is not, so the two convey complementary information.

### 0.11 Feature importance and ablation analysis

Global feature importance was visualised using a beeswarm plot^52^ of SHAP (SHapley Additive exPlanations) LinearExplainer values^53^ computed from the Logistic Regression classifier.

The Leave-one-feature-out (LOFO) ablation was performed by iteratively retraining the selected model with each feature removed and computing the resulting change in cross-validated MCC (ΔMCC) over the same grouped folds. A complementary feature-family ablation analysis refit the selected model on nested subsets of the selected features: conservation and sequence-specific descriptors alone (*n* = 5), structural descriptors alone (*n* = 6), the complete set (*n* = 11), and the complete set with the stability (*n* = 1) or conservation (*n* = 4) features withheld. This isolates how much the structural descriptors add beyond sequence conservation. Each subset was evaluated by pooled out-of-fold grouped cross-validation and on the held-out test set, with bootstrap 95% confidence intervals (2000 resamples).

### 0.12 Independent validation and benchmarking

UshEffect-3D was additionally evaluated on the 78 ACMG-classified ^36^ pathogenic variants held out from all training and evaluation steps. On this validation set, we primarily focus on sensitivity as it contains only pathogenic variants due to which other metrics can be misleading.

We additionally benchmarked UshEffect-3D against six general-purpose variant effect predictors on the held-out test set including PolyPhen-2^54^, SNAP2^55^, VESPAl^56^, AlphaMissense^57^, ESM-1b^58^, and CPT-1^59^, all evaluated on the same 110 variants. Each tool’s continuous score was thresholded at its published cutoff where one is defined. CPT-1 publishes no decision cutoff^59^, so we applied the conventional 0.50 to its output, which is the modelled probability of the functionally abnormal class.

As point estimates alone cannot establish superiority on a test set of this size, every metric is reported with a bootstrap 95% confidence interval (2000 case resamples). The difference between UshEffect-3D and each predictor was estimated as a paired difference, with both predictors scored on identical resampled variants. A difference was treated as statistically resolvable only where its 95% confidence interval excluded zero.

## Results

### 1 USH2A variant burden and domain architecture define the structural problem

Clinically observed mutations occur throughout Usherin, but they are not evenly distributed across its domain landscape (Figure 1B; Supplementary Table S1). Adjusting for domain size, classified variants are reported most densely in the Laminin N-terminal (17.8 variants per 100 residues) and Laminin G-like domains (14.0 per 100 residues), followed by a comparatively modest distribution within the Laminin EGF-like (12.1 per 100 residues) and Fibronectin type III (11.6 per 100 residues) domains.

Pathogenic variants are significantly enriched in the Laminin N-terminal (37 of 44 variants; odds ratio 2.94, 95% CI 1.29–6.72, *q* = 0.013) and Laminin G-like domains (43 of 52; OR 2.68, 95% CI 1.28–5.60, *q* = 0.013), and significantly depleted across the Fibronectin type III repeats (226 of 372; OR 0.57, 95% CI 0.40–0.81, *q* = 0.009). This depletion is proportional, not absolute. The Fibronectin type III repeats span 3213 of the 5202 residues and contain 226 of the total 407 pathogenic variants, more than all other families combined, even though pathogenic variants are under-represented within them relative to the rest of the protein. The Laminin EGF-like domains, however, trend towards enrichment without reaching significance (OR 1.54, 95% CI 0.86–2.75, *q* = 0.21). This shows that domain identity alone may not be sufficient to prioritise risk, motivating a residue-level approach.

### 2 AlphaFold2-derived domain models recover the expected fold architecture

Across all 47 domain instances spanning the four constituent fold types, the AlphaFold2-derived domain models recovered the expected fold architecture (Figure 2A). Across the 4389 modelled residues of these instances, of which 3931 aligned to a template, the residue-weighted median TM-score was 0.71, comfortably above the 0.5 threshold for correct fold assignment^34^, and 43 of 47 domain instances (91%) scored *≥* 0.5 (Figure 2B). Additionally, agreement was consistent within each domain family (Table 1). Fibronectin type III domains, the dominant tandemly repeated fold, clustered tightly (median TM 0.71, interquartile range 0.66–0.74, *n* = 34; median C*α* RMSD 2.45 Å), as did the Laminin EGF-like domains (median TM 0.72, IQR 0.67–0.80, *n* = 10; RMSD 1.78 Å). The two Laminin G-like domains showed the closest correspondence (median TM 0.86; RMSD 2.20 Å), and the single Laminin N-terminal domain recovered its fold at TM 0.66 (RMSD 2.47 Å).

**Figure 2:**
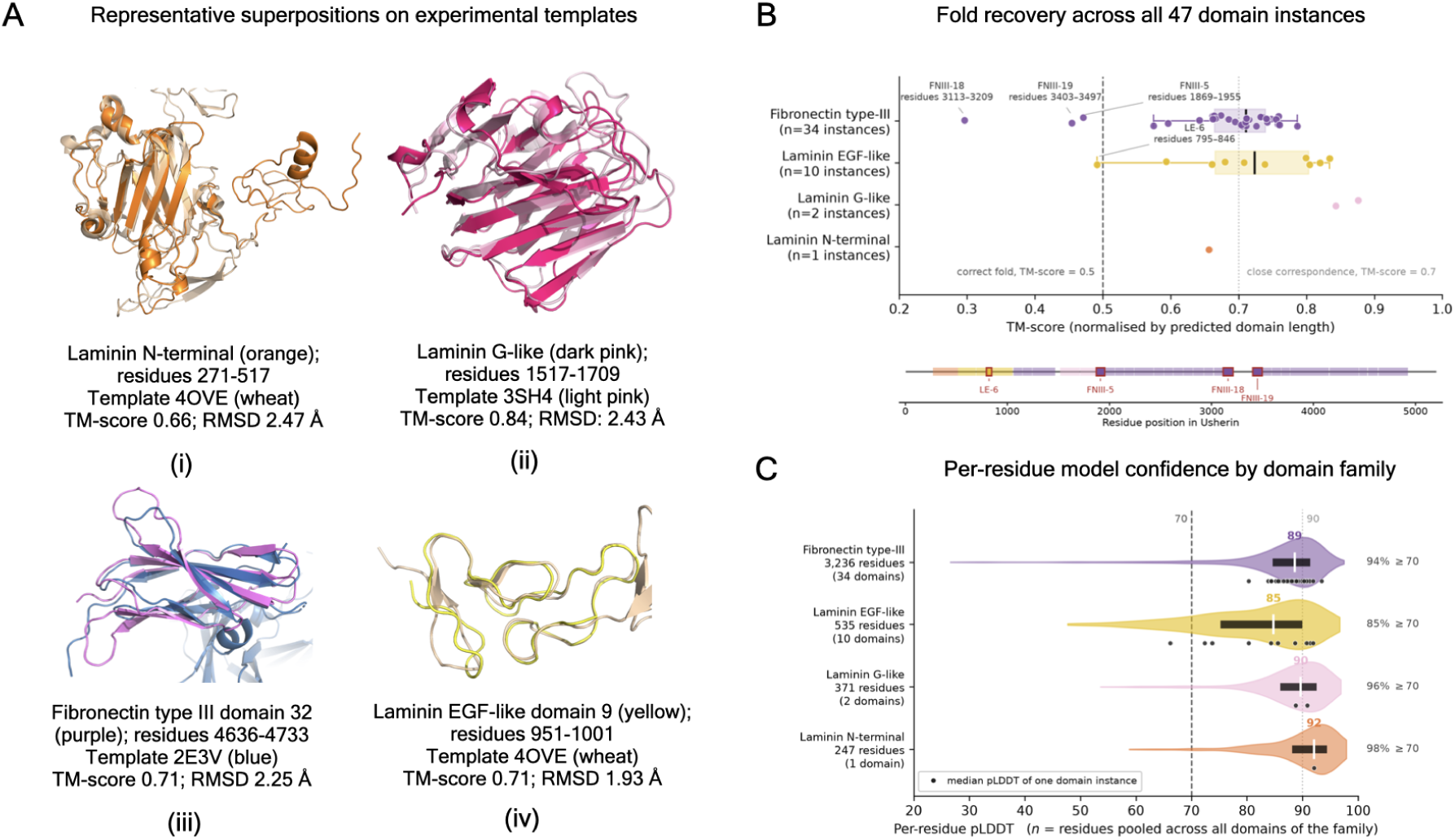
Fold concordance and model confidence of the AlphaFold2-derived Usherin domains, assessed against representative experimental structures of the same fold families. **(A)** Representative superpositions of AlphaFold2-predicted domains (solid colour) on their experimental templates (pale), each showing the instance whose TM-score lies closest to its family median: (i) Laminin N-terminal, residues 271–517 (orange) on 4OVE (wheat); (ii) Laminin G-like 1, residues 1517–1709 (dark pink) on 3SH4 (light pink); (iii) Fibronectin type III 32, residues 4636–4733 (purple) on 2E3V (blue); (iv) Laminin EGF-like 9, residues 951–1001 (yellow) on 4OVE (wheat). Each is annotated with its TM-score and C*α* RMSD. Superpositions reproduce the TM-align alignment in PyMOL version 2.5.0^28^. **(B)** TM-scores for all 47 domain instances, normalised by predicted-domain length and grouped by family. Each point is one instance. Boxes span the interquartile range with the median marked, and are drawn only for families with at least five instances, so the Laminin G-like (*n* = 2) and Laminin N-terminal (*n* = 1) families are shown as individual points and no distribution is implied from so few observations. The dashed line marks the TM=0.5 threshold for recovery of the correct fold and the dotted line TM=0.7 for close structural correspondence. The four instances falling below 0.5 are labelled with their domain family and residue range. The strip beneath locates all 47 instances along the 5202-residue sequence in family colours, with those four outlined in red, showing that the sub-threshold cases are dispersed rather than clustered in one region of the protein. **(C)** Distribution of per-residue AlphaFold2 pLDDT within each family, shown as violins with the interquartile range and median overlaid. Here *n* is the number of residues pooled across all domains of the family, and the number of contributing domains is given separately. Each black point is the median pLDDT of a single domain instance, which distinguishes low-confidence residues from a whole domain modelled at low confidence. One Laminin EGF-like instance has a median below 70 on this basis. The dashed and dotted lines mark the conventional pLDDT 70 and 90 confidence thresholds respectively, and the percentage of residues at pLDDT *≥*70 is given at the right of each row.

**Table 1:** Per-domain-family fold concordance and model confidence for the AlphaFold2-derived Usherin domains. TM-scores (normalised by predicted-domain length), C*α* RMSD and sequence identity were obtained by TM-align against a representative experimental structure of each fold family; pLDDT is the AlphaFold2 per-residue confidence, summarised over the modelled residues of each family. Low sequence identity alongside high TM-scores indicates that the observed agreement reflects genuine structural correspondence. Per-instance values for all 47 domains are provided in Supplementary Table S2.

| Domain family | Template | *n* | Residues | Median TM (IQR) | Min TM | Median RMSD (Å) | Median seq. ID (%) | Median pLDDT (IQR) | Res. $\geq 70$ (%) | Res. $\geq 90$ (%) |
| --- | --- | --- | --- | --- | --- | --- | --- | --- | --- | --- |
| Fibronectin type III | 2E3V | 34 | 3236 | 0.71 (0.66-0.74) | 0.30 | 2.45 | 17.8 | 88.6 (84.6-91.3) | 94.4 | 38.0 |
| Laminin EGF-like | 4OVE | 10 | 535 | 0.72 (0.67-0.80) | 0.49 | 1.78 | 35.7 | 84.8 (75.2-90.0) | 85.2 | 24.7 |
| Laminin G-like | 3SH4 | 2 | 371 | 0.86 (0.85-0.87) | 0.84 | 2.20 | 22.1 | 89.7 (85.9-92.5) | 96.0 | 46.6 |
| Laminin N-terminal | 4OVE | 1 | 247 | 0.66 (-) | 0.66 | 2.47 | 26.8 | 92.0 (88.0-94.4) | 98.0 | 64.8 |
| All families |  | 47 | 4389 | 0.71 | 0.30 | - | 19.8 | 88.5 (84.1-91.5) | 93.6 | 38.6 |

Four instances fell below the fold-recovery threshold (Figure 2B): three Fibronectin type III repeats – FNIII-5 (residues 1869–1955, TM 0.47), FNIII-18 (3113–3209, TM 0.30) and FNIII-19 (3403–3497, TM 0.45) – and one Laminin EGF-like domain, LE-6 (795–846, TM 0.49). As the sequence strip in Figure 2B shows, these are dispersed across the extracellular domains rather than concentrated in one region. Alignment was incomplete in each case wherein 54 of 87 modelled residues aligned for FNIII-5 (62%), 41 of 97 for FNIII-18 (42%) and 54 of 95 for FNIII-19 (57%). The quality of the aligned cores differed, however. FNIII-19 superposed to 2.01 Å, better than the family median of 2.45 Å, so its low TM-score is attributable almost entirely to the extent of alignment rather than to the fit of the aligned region. FNIII-18, by contrast, combined the poorest alignment with the poorest superposition (3.16 Å), and FNIII-5 was intermediate (2.59 Å). For LE-6 however, alignment was near-complete with 48 of 52 residues (92%), which implies that the low score does not arise from partial alignment but from a relatively poorer superposition. At 2.9 Å C*α* RMSD, it is the least well-fitting of the ten Laminin EGF-like repeats.

Per-residue model confidence was correspondingly high (Figure 2C; Table 1). Across the four families the median pLDDT was 88.5 (interquartile range 84.1–91.5), with 93.6% of residues at pLDDT *≥*70 and 38.6% at *≥*90. Confidence was highest for the Laminin N-terminal domain (median 92.0) and lowest within the Laminin EGF-like family (median pLDDT 84.8; 85.2% of residues *≥*70). However, resolving confidence to individual instances shows that this shortfall is concentrated in a single domain - LE-6 has a median pLDDT of 66, the only instance of the 47 to fall below 70, while every other instance exceeds that threshold (Figure 2C). LE-6 is also the Laminin EGF-like instance that failed the fold-recovery threshold (TM 0.49), so the two independent measures converge on the same domain.

Taken together, these results establish that the predicted Usherin domains are, with a few localised exceptions, accurate at the fold level. Fold-level validation was, however, restricted to the four extracellular domain families for which experimental templates were available. This delimits the regions in which the structure-derived features underpinning the downstream model are least reliable.

### 3 A biologically interpretable model supports accurate variant classification

#### 3.1 A simple linear model demonstrated the most stable cross-validated performance

We evaluated eleven machine learning architectures by nested cross-validation, grouping variants by residue position to avoid data leakage between splits. Mean nested-CV MCC was consistently uniform across architectures, spanning between 0.73 to 0.77 (Figure 3A; Supplementary Table S3). Discrimination is therefore effectively saturated by the selected feature set, and the choice between architectures cannot be resolved on mean performance alone.

**Figure 3:**
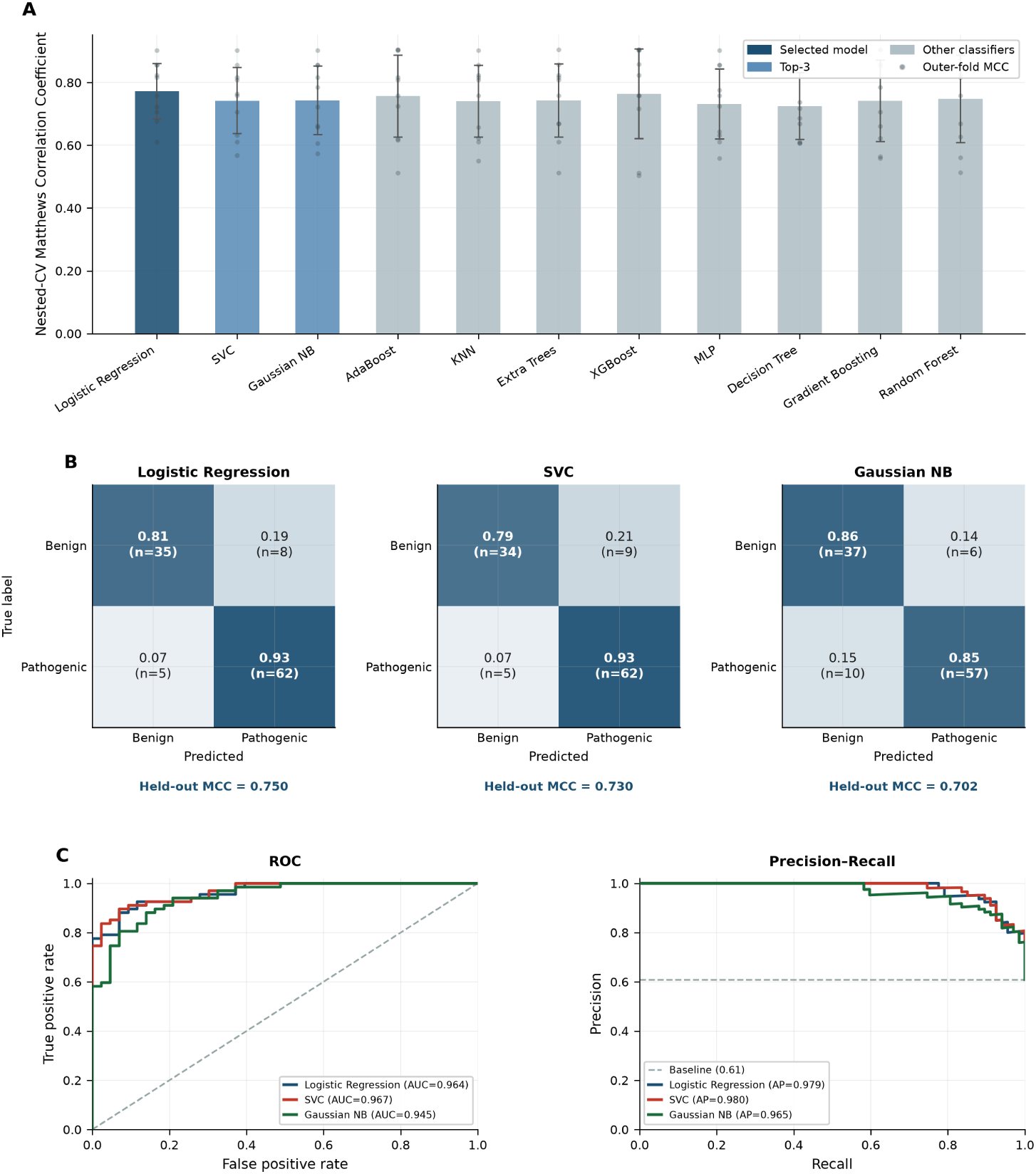
Training and evaluation summary of all classifiers to inform model selection. **(A)** Nested position-grouped cross-validation Matthews Correlation Coefficient (MCC) for 11 classifiers. Bar height represents the mean across the ten outer folds, error bars represent the standard deviation, and grey points are the individual outer-fold scores. Classifiers are ordered by the selection criterion (mean MCC minus its standard deviation). The selected model is highlighted in dark blue and the remaining top-three models in mid blue. Mean performance is closely comparable across architectures, so the ranking is driven largely by fold-to-fold consistency. **(B)** Confusion matrices for the three top-ranked classifiers on the held-out test set, showing the row-normalised proportion and count of true benign and pathogenic predictions. Among the models that displayed the most stable cross-validation performance, Logistic Regression achieved the highest held-out MCC (0.750), followed by Support Vector Classifier (0.730) and Gaussian Naive Bayes (0.702). Gaussian Naive Bayes retained the most benign variants (37 of 43) but missed the most pathogenic ones (10 of 67). **(C)** Receiver operating characteristic (ROC) and precision-recall curves for the top-three models on the held-out test set. The top three models achieved near-identical discrimination with comparable AUROC and Average Precision.

Models were consequently ranked by mean nested-CV MCC minus its standard deviation across the ten outer folds (Supplementary Table S3), which rewards consistency between folds as well as average accuracy. Logistic Regression ranked first on this criterion (0.68), combining the highest mean MCC (0.77 *±* 0.09) with the lowest fold-to-fold variability of any classifier. The Support Vector Classifier (0.64) and Gaussian Naive Bayes (0.63) models closely followed. Notably, tree-based, ensemble methods such as XGBoost (0.76 *±* 0.14) and Random Forest (0.75 *±* 0.14) achieved comparable mean MCC but with slightly greater instability. The failure of ensemble methods to gain from their capacity to model feature interactions suggests that the selected features separate the classes largely additively, thus favouring a simple, linear model.

On the held-out set (*n* = 110), Logistic Regression achieved an MCC of 0.75 (Table 2), closely tracking its cross-validated performance, and reached a sensitivity of 0.76 on the independent ACMG validation set. The Support Vector Classifier performed near-identically (MCC 0.73), while Gaussian Naive Bayes (MCC 0.70) traded sensitivity for specificity, correctly identifying 37 benign variants but missing 10 pathogenic ones (Figure 3B). The higher specificity of the Gaussian Naive Bayes model was surprising considering the training set contained more pathogenic than benign variants.

**Table 2:** Performance summary of the selected model (Logistic Regression - UshEffect-3D) across nested position-grouped cross-validation (10 outer folds, mean *±* SD), the group-aware held-out test set, and the independent ACMG validation set. MCC, accuracy, balanced accuracy, F1, precision and specificity are not reported (–) for the ACMG set, which contains only pathogenic variants.

| Dataset | MCC | Accuracy | BAcc | F1 | Precision | Sensitivity | Specificity |
| --- | --- | --- | --- | --- | --- | --- | --- |
| Nested CV | $0.77 \pm 0.09$ | $0.89 \pm 0.04$ | $0.88 \pm 0.05$ | $0.91 \pm 0.03$ | $0.91 \pm 0.06$ | $0.92 \pm 0.05$ | $0.84 \pm 0.11$ |
| Test | 0.75 | 0.88 | 0.87 | 0.91 | 0.89 | 0.93 | 0.81 |
| ACMG | – | – | – | – | – | 0.76 | – |

The ROC curves (Figure 3C) give the probability that a randomly chosen pathogenic variant is scored above a randomly chosen benign one at different thresholds. Discrimination was high for all three classifiers against the 0.5 random-classifier baseline (AUROC 0.964, 0.967 and 0.945 for Logistic Regression, the Support Vector Classifier and Gaussian Naive Bayes respectively), with negligible differences between them. Given that our dataset contained more pathogenic than benign variants, the PR curve allows us to evaluate the trade-off between precision and recall. All three classifiers achieved high Average Precision against a prevalence baseline of 0.609 - 0.979, 0.980 and 0.965 for Logistic Regression, the Support Vector Classifier and Gaussian Naive Bayes respectively (Figure 3C), indicating that each model recovers most pathogenic variants while keeping false positives low, with negligible differences between them.

#### 3.2 Evolutionary conservation dominates model predictions

To establish the importance of individual features to model predictions, SHAP (SHapley Additive exPlanations) values were computed across the training variants (Figure 4A; Supplementary Table S6), where positive SHAP values indicate a contribution toward the pathogenic class (class 1) and negative values toward the benign class (class 0)^60^. Evolutionary constraint features dominated model output by magnitude and spread - Shannon entropy (`Demask_entropy`, mean *|*SHAP*|* = 0.131) and the log of variant frequency (`Demask_log2f_var`, 0.130) exhibited the widest SHAP distributions, followed by `delta_PSIC` (0.107) and `Consurf_score` (0.098), indicating that these four sequence-based features exert the greatest overall influence on individual predictions. `DDmut` ΔΔ*G* ranked fifth (0.076), retaining a subtle contribution.

**Figure 4:**
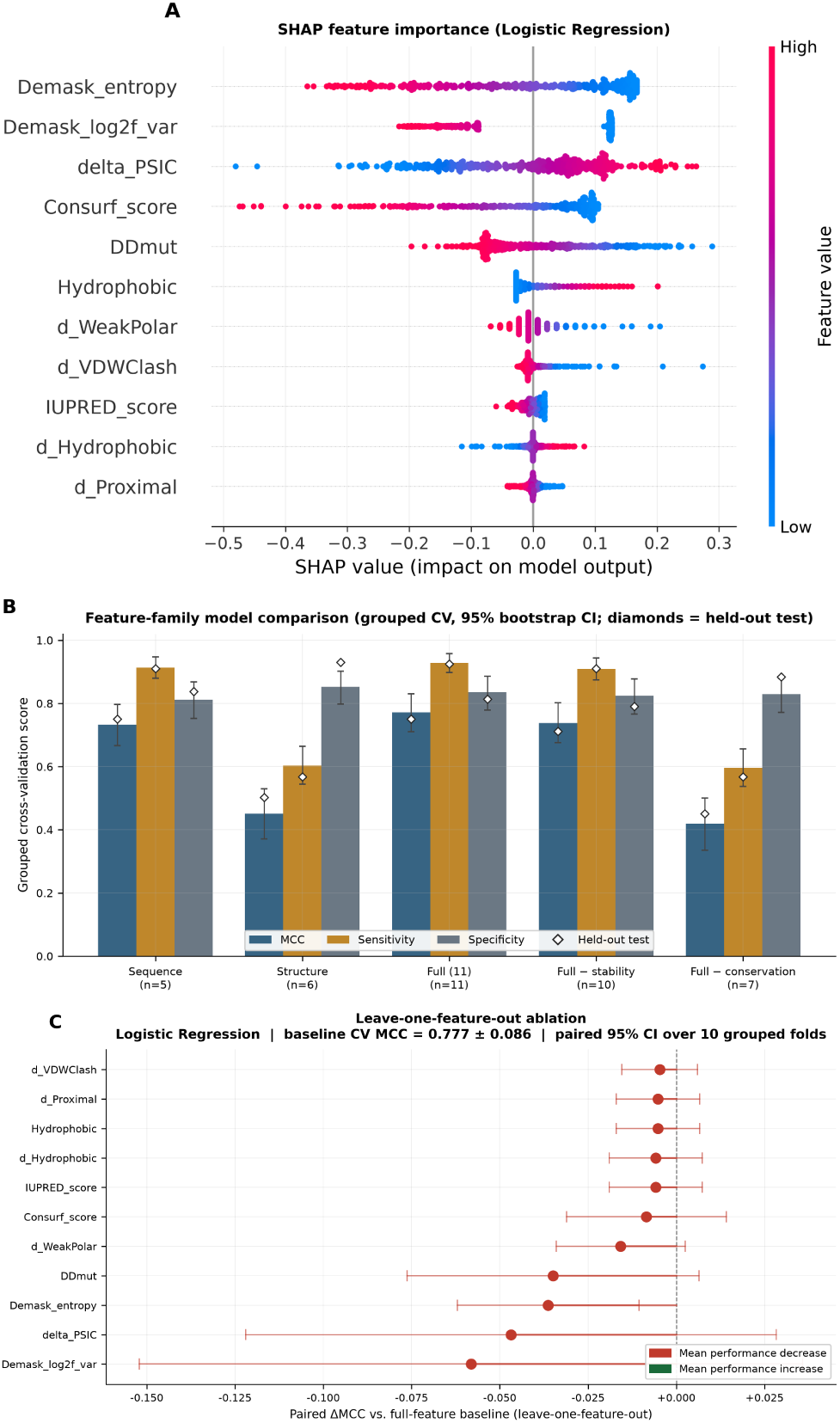
Feature importance and ablation analyses for UshEffect-3D. **(A)** SHAP global feature importance for the selected Logistic Regression classifier, computed with LinearExplainer, showing the distribution of SHAP values for each of the eleven selected features across the training variants. Each point represents a single variant, coloured by its relative feature value (red = high, blue = low). Positive values indicate a contribution toward the pathogenic class and negative values toward the benign class. Features are ordered by mean absolute SHAP value, with Demask_entropy and Demask_log2f_var the most influential, followed by delta_PSIC, Consurf_score and DDmut. The six remaining structural-environment and disorder descriptors show narrow distributions concentrated near zero. **(B)** Feature-family model comparison. The selected model was refit on conservation and sequence descriptors alone (*n* = 5), structural descriptors alone (*n* = 6), the complete set (*n* = 11), and the complete set with the stability or conservation features withheld. Bars show grouped cross-validation MCC, sensitivity and specificity with bootstrap 95% confidence intervals (2000 resamples); diamonds mark the corresponding single held-out test score. Conservation alone recovers essentially the full performance, whereas structure alone does not. **(C)** Leave-one-featureout (LOFO) ablation for the selected classifier (baseline CV MCC = 0.777*±*0.086), showing the mean ΔMCC upon individual removal of each feature. Intervals are paired 95% confidence intervals on the within-fold difference from the full model across the ten grouped folds. Every feature produced a small, negative mean ΔMCC, the largest being Demask_log2f_var (*−*0.058) and delta_PSIC (*−*0.047). No individual feature seems to be independently indispensable.

The direction of each contribution is biologically coherent. Low entropy and low variant frequency both push predictions toward pathogenicity, consistent with rare substitutions at evolutionarily constrained positions being more likely to disrupt function^24,39^. ConSurf behaves equivalently, with slower rates of evolution favouring a pathogenic call^38^. Higher `delta_PSIC`, which reflects a greater divergence between wild-type and mutant residue propensities, like-wise drives predictions toward pathogenicity^41^. For `DDmut`, destabilising substitutions (more negative ΔΔ*G*) were shifted toward pathogenic predictions indicating the loss of local fold integrity as a likely disease mechanism^45^. The six remaining structural-environment descriptors — wild-type hydrophobic contacts (`Hydrophobic`; 0.031), change in weak-polar contacts upon mutation (`d_WeakPolar`; 0.023), change in Van der Waals clash upon mutation (`d_VDWClash`; 0.014), hydrophobic (`d_Hydrophobic`; 0.011) and proximal (`d_Proximal`; 0.007) contacts, and the IUPred3 disorder score (`IUPRED_score`; 0.012) — displayed narrow SHAP distributions concentrated near zero, indicating modest and largely uniform contributions compared to conservation and stability features.

#### 3.3 Grouped feature ablation uncovers redundancy among structural descriptors

To assess the collective contribution of functionally related feature groups, the selected model was refit on subsets of the eleven features and evaluated using the grouped cross-validation strategy (without retuning) (Figure 4B; Table S7). The complete feature set achieved a pooled out-of-fold MCC of 0.77 (95% CI 0.71–0.83). Conservation and sequence descriptors alone recovered almost all of this signal (MCC 0.73, 95% CI 0.67–0.80), whereas the structural descriptors alone performed substantially worse (MCC 0.45, 95% CI 0.37–0.53), establishing that evolutionary information carries the bulk of the discriminative power. Withholding the single stability feature (`DDmut`) lowered cross-validated MCC by 0.035, but with enough fold-to-fold variability that no reduction at all remains consistent with the data (95% CI *−*0.076 to +0.006). The difference between the full model and conservation features alone is likewise indistinguishable from nothing (*−*0.042, 95% CI *−*0.085 to +0.001).

The evidence therefore supports that evolutionary features provide the dominant predictive signal, while the structural descriptors supply at most a small complementary contribution that this dataset cannot resolve from zero. Notably, the two are indistinguishable on the held-out set as well, where conservation alone and the full feature set achieved MCCs of 0.751 and 0.750 respectively (Figure 4B). We therefore do not claim that adding structural features improves overall discrimination. Their contribution, if real, is small relative to the between-fold variability of this dataset.

At the individual feature level, a leave-one-feature-out (LOFO) ablation was performed against a full-feature baseline of 0.78 *±* 0.09, the mean of the ten per-fold MCCs (Figure 4C). Every feature produced a very small, negative mean ΔMCC on removal, the largest being `Demask_log2f_var` (*−*0.058), `delta_PSIC` (*−*0.047), `Demask_entropy` (*−*0.036) and `DDmut` (*−*0.035). The remaining seven each changed MCC by less than 0.016. No individual feature is therefore independently indispensable to model performance, consistent with the additive feature contribution leveraged by the linear model.

### 4 UshEffect-3D performs comparably to general-purpose predictors with a better-calibrated decision threshold

A key question we tried to answer is whether gene-specific models offer clear advantages over established, general-purpose tools. In a head-to-head comparison on the group-aware held-out test set, UshEffect-3D achieved the highest MCC (0.75, 95% CI 0.62–0.87) and balanced accuracy (0.87, 0.80–0.93) of the seven predictors evaluated (Table 3), ahead of PolyPhen-2^54^ (MCC 0.66), VESPAl^56^ and ESM-1b^58^ (0.65), AlphaMissense^57^ (0.64) and CPT-1^59^ (0.63), with SNAP2^55^ considerably worse (0.26). However, in terms of threshold-free ranking ability described by AUROC, all tools except for SNAP2^55^ scored higher than 0.9 (Table 3).

**Table 3:**
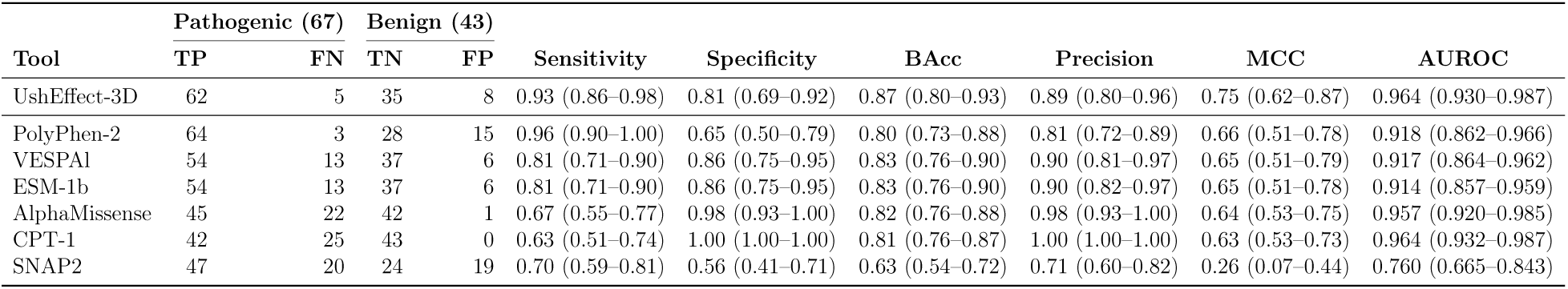
Benchmarking UshEffect-3D against general-purpose variant effect predictors on the group-aware held-out test set (110 variants; 67 pathogenic, 43 benign), sorted by MCC. Every metric is given as the point estimate with a bootstrap 95% confidence interval (2000 resamples). Paired differences between UshEffect-3D and each competing predictor are given in Supplementary Table S5.

Given the imbalanced composition of the test set (67 pathogenic, 43 benign), we looked into the distribution of errors to ascertain whether general purpose predictors were robust to class imbalance. PolyPhen-2 misclassified 15 of the 43 benign variants, a bias toward pathogenic calls that inflates sensitivity (0.96) at the cost of specificity (0.65). AlphaMissense misclassified a single benign variant and CPT-1 misclassified none of the benigns but they missed 22 and 25 of the 67 pathogenic variants respectively (sensitivity 0.67 and 0.63), thus leaning towards a more conservative prediction profile. UshEffect-3D sits between these failure modes with 8 false positives and 5 false negatives (sensitivity 0.93, specificity 0.81). The tools differ far more in where their errors fall than in how well they rank variants, which could be explained by their poorly calibrated proteome-wide decision thresholds rather than in their discriminative power.

Point estimates alone do not establish superiority on a test set of this size. Estimated as paired differences, with both predictors scored on identical resampled variants, UshEffect-3D’s advantage in MCC is statistically resolvable only over SNAP2 (ΔMCC = +0.49, 95% CI +0.25 to +0.72). Against PolyPhen-2, VESPAl, ESM-1b, AlphaMissense and CPT-1 the margins are small and every interval spans zero (+0.09 to +0.12; Supplementary Table S5). The same comparison on AUROC places UshEffect-3D marginally ahead of VESPAl and ESM-1b (ΔAUROC +0.046, 95% CI +0.006 to +0.091, and +0.050, +0.013 to +0.093) and indistinguishable from PolyPhen-2 (+0.046, *−*0.001 to +0.098), AlphaMissense (+0.007, *−*0.030 to +0.043) and CPT-1 (*−*0.001, *−*0.025 to +0.020). Therefore, on these 110 variants UshEffect-3D is best described as performing comparably to the leading general-purpose predictors, differing from them in the placement of its decision threshold rather than in demonstrable discriminative power.

### 5 Reclassification of ClinVar VUS highlights clinically relevant variants across the protein

Applying this framework to the 2639 *USH2A* VUS currently in ClinVar, our model prioritised 37.2% (*n* = 981) as likely pathogenic and 62.8% (*n* = 1658) as likely benign (Figure 5A). These predictions are distributed across the protein with no clear hotspots (Figure 5B), and VUS density is near uniform across domain types once domain size is accounted for, ranging from 49.7 to 53.9 VUS per 100 residues (Figure 5C). The predicted probabilities themselves occupy a narrow band, spanning 0.21 to 0.75 with 92.5% of VUS scoring between 0.3 and 0.7, so few variants are assigned to either class with very high confidence.

**Figure 5:**
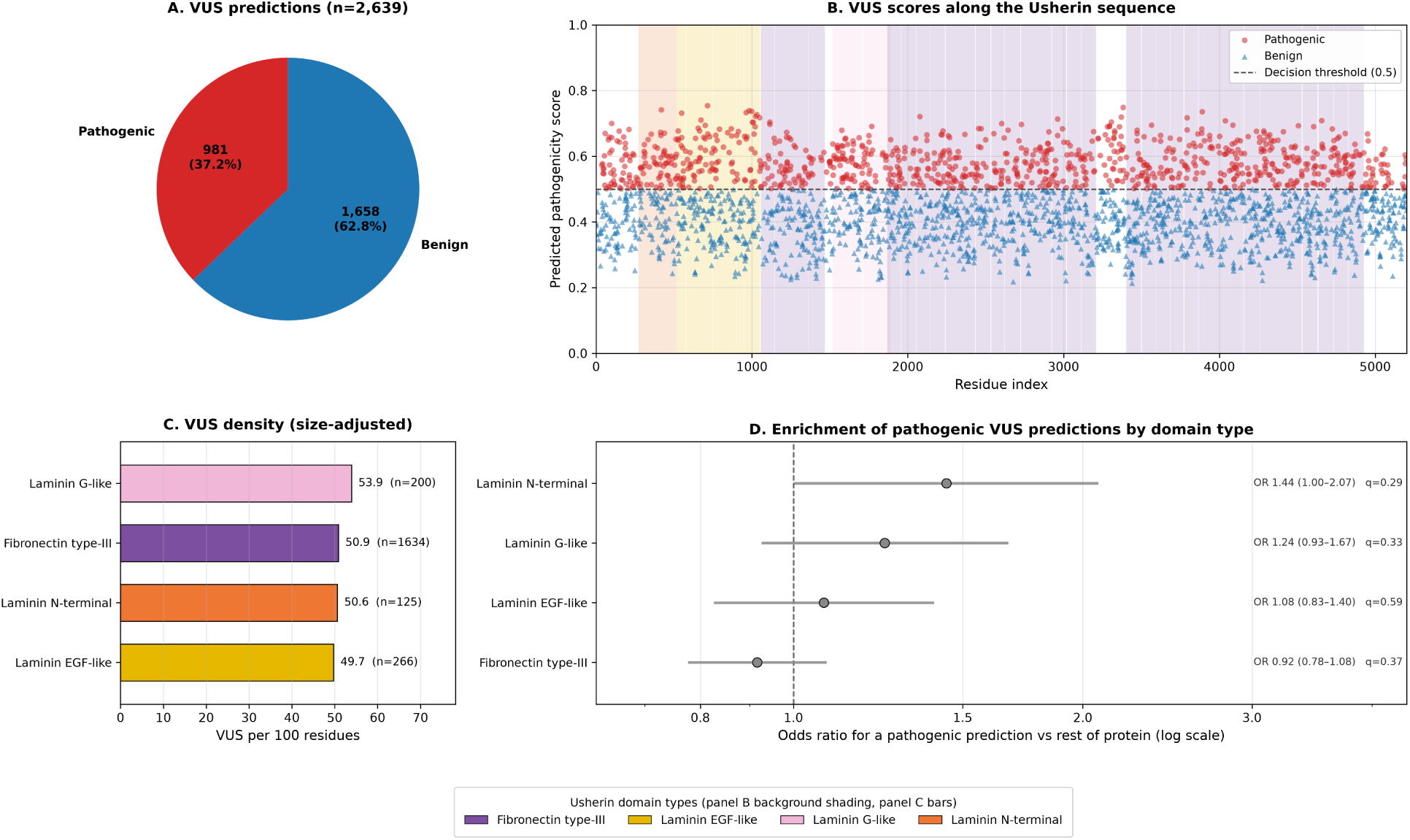
UshEffect-3D predictions across *USH2A* variants of uncertain significance. **(A)** Pie chart showing pathogenicity predictions across all 2639 *USH2A* VUS, with 62.8% predicted benign and 37.2% predicted pathogenic. **(B)** Scatter plot showing predicted pathogenicity scores against residue position across the full length of the USH2A protein. Each point represents a single VUS, coloured and shaped by predicted class (red circles = pathogenic, blue triangles = benign); the dashed line marks the 0.5 decision threshold. Background shading denotes annotated protein domains, as indicated in the legend. Pathogenic predictions are distributed across the entire protein length, with no obvious positional clustering. **(C)** VUS density by domain type, expressed as the number of VUS per 100 residues of that domain type — the quantity that adjusts for domain size (see Methods). Density is near uniform across the four domain families (49.7–53.9 per 100 residues). **(D)** Enrichment of pathogenic-predicted VUS by domain type, shown as odds ratios with 95% confidence intervals for a pathogenic prediction within each domain type against the remainder of the protein (Fisher’s exact test, Benjamini-Hochberg corrected across domain types). All intervals span an odds ratio of 1 and no domain type reaches *q <* 0.05, so the predicted pathogenic VUS show no domain-level enrichment. The PDZ binding motif and transmembrane segment are excluded throughout as they are too small to support inference.

The proportion of VUS predicted pathogenic was seemingly highest in the Laminin N-terminal (45.6%) and Laminin G-like domains (42.0%) against a protein-wide prediction rate of 37.2%. However, when comparing predicted pathogenic counts against benign counts within each domain type to the remainder of the protein, no domain type reached significance after correction for multiple testing. The strongest signal, the Laminin N-terminal domain, gave an odds ratio of 1.44 (95% CI 1.00–2.07) with a corrected *q* of 0.29, followed by the Laminin G-like domains (OR 1.24, 95% CI 0.93–1.67, *q* = 0.33), while the Fibronectin type III repeats moved towards depletion without reaching significance (OR 0.92, 95% CI 0.78– 1.08, *q* = 0.37; Figure 5D). The predicted pathogenic VUS are therefore distributed across the Usherin domain landscape in a manner statistically indistinguishable from uniform, and we do not claim domain-level enrichment among them.

## Discussion

Usherin is among the largest proteins in the human proteome, spanning 5202 residues across four major extracellular domain families^7,9^, with no known, full-length experimental structure in public databases^12^. The sheer size of this protein along with the vast number of VUS associated with it makes variant interpretation through functional assays impractical^61^. We therefore generated AlphaFold2 models of all 47 annotated domains, validated their folds against experimental templates of the same families, derived residue-level structural descriptors from them, and assessed whether these descriptors improve missense variant classification over sequence-based features alone.

### 5.1 Structural features add no discrimination beyond evolutionary conservation

SHAP analysis identified evolutionary constraint features, specifically Shannon entropy (`Demask_entropy`)^39^ and variant frequency with respect to residue position (`Demask_log2f_var`), both derived from DeMaSk^24^, the divergence in residue propensity on substitution (`delta_PSIC`^41,42^) and rate of evolution (`Consurf_score`^38^), as the dominant contributors to model predictions. This is biologically coherent, as positions under strong purifying selection are by definition those where amino acid identity is functionally critical^62^. The feature-family ablation analysis confirmed it quantitatively as the sequence-based features alone recovered a cross-validated MCC of 0.73 against 0.77 for the complete feature set, whereas structural features alone reached only 0.45. The held-out set gives the same answer independently, where conservation features alone achieved an MCC of 0.751. Thus, the structural descriptors seem to add nothing detectable beyond what sequence conservation already provides.

Two explanations are plausible, and they are not mutually exclusive. Evolutionary conservation metrics are themselves partly a readout of structural importance, so buried, tightly packed and structurally critical positions are already flagged as conserved ^63,64^. Alternatively, the current set of structural features may lack critical inter-domain context. Features were deliberately calculated within single excised domains to avoid spurious contacts arising from unreliable inter-domain placement, which necessarily excludes the interface and homomerisation geometry through which several *USH2A* variants are unaccounted for ^65^.

The ablation also illustrates why individual and grouped feature analyses must be read together. Every one of the eleven features produced a minor negative mean ΔMCC on removal (Figure 4C), yet withholding the conservation family as a group cost 0.35 MCC (Table **??**). This reinstates that features within each group carry complementary information that only become apparent when removed as a whole. `Consurf_score` and `Demask_entropy` both summarise how strongly a position resists substitution, differing only in whether the score is estimated from evolutionary rate or from alignment column diversity ^24,38,39^. `delta_PSIC` and `Demask_log2f_var` instead incorporate the identity of the substituted residue^24,41,42^, and it is their pairing with the position-level measures that yields the largest gains. The conservation feature set therefore encodes two distinct axes of evolutionary information, and any single descriptor can be withheld because at least one measure of its axis remains, while withholding the family removes both axes at once.

Critically, it is important to clarify that ablation studies and SHAP analyses only tell us what the model considers important^66^, not that certain features or predicted structures themselves are completely redundant. Moreover, the lack of signal from structural descriptors cannot be attributed to poorly predicted domains as the fold validation analysis showed that 43 of 47 domain instances recovered their fold at TM *≥* 0.5, and 93.6% of residues rest above the expected pLDDT confidence threshold. Thus, in the case of Usherin, the application of interpretable machine learning techniques do not make up for the lack of functional validation required to understand the mechanisms of variation.

### 5.2 A gene-specific model does not offer a clear advantage over general-purpose predictors

The selected Logistic Regression classifier achieved consistent performance on unseen data (Table 2), with held-out metrics closely tracking their cross-validated counterparts. In the context of treatment eligibility assessment, the consequences of classification errors are serious - a false positive risks enrolling a patient in an expensive treatment program on an incorrect basis, while a false negative risks denying an affected patient access to a potentially sight-preserving intervention. UshEffect-3D’s sensitivity of 0.93 limits the probability that a clinically significant variant is overlooked, though its comparatively lower specificity of 0.81 means roughly one in five benign variants may be flagged as pathogenic. Thus, we urge clinicians to use these predictions for prioritising variants for review rather than for standalone classification.

The general-purpose predictors differ from our model far more in where their errors fall than in how well they rank variants. PolyPhen-2 achieved nominally competitive sensitivity (0.96) at the cost of much lower specificity (0.65). At the opposite extreme, AlphaMissense and CPT-1 misclassified one benign variant and none respectively, but recovered only 67% and 63% of pathogenic variants. Since threshold-free ranking ability was comparable across these tools, the gene-specific advantage is best understood as a decision threshold calibrated to the class balance of this gene rather than as evidence that our model captures determinants of pathogenicity the others miss.

A methodological point deserves emphasis, as it bears on any variant effect predictor built for a single protein. As multiple substitutions frequently occur at the same residue, splitting variants without regard to position allows the same residue to inform both training and evaluation, and the structural and conservation features used here are largely position-determined. Enforcing group-aware splits and nested, position-grouped cross-validation throughout yields more conservative performance estimates than a conventional stratified split would, and we regard this as the appropriate standard for gene-specific models built on variant sets. It also explains why the eleven classifier architectures we compared proved statistically indistinguishable from one another - once positional leakage is removed, the discriminative ceiling is set by the feature set rather than by the choice of architecture.

### 5.3 Reclassification of ClinVar VUS expands the landscape of pathogenic variants

UshEffect-3D prioritised 37.2% of *USH2A* ClinVar VUS as likely pathogenic. These predictions were distributed across Usherin without detectable domain-level enrichment. The Laminin N-terminal and Laminin G-like domains carried the highest predicted pathogenic proportions (45.6% and 42.0% against a protein-wide 37.2%), consistent with the established roles of these domains in extracellular matrix binding and protein-protein interactions within the USH2 complex^9,65^. However, the corresponding odds ratios among VUS were much closer to unity (1.44 and 1.24) and did not reach significance, so this correspondence should be regarded as suggestive rather than established.

Prioritisation of the 981 likely pathogenic VUS nonetheless represents a tractable target for directed functional validation and, where evidence is sufficient, upgrading clinical classification under ACMG/AMP criteria^36^. Given that gene editing and antisense oligonucleotide strategies targeting *USH2A* are under active investigation^67^, variant resolution at this scale could meaningfully expand the pool of patients eligible for trials and approved therapies in the future. We note, however, that the predicted probabilities from UshEffect-3D’s linear model occupy a comparatively narrow range, so the ranking within the prioritised set should be treated as indicative and used to order variants for review rather than as a calibrated measure of confidence in any individual call.

### 5.4 Limitations and future directions

Several limitations of the present study warrant consideration. The model was developed and evaluated exclusively on *USH2A* missense variants. Its performance on other variant classes including in-frame indels, synonymous variants with splicing consequences, or deep intronic variants, is unknown and should not be assumed. The structural features used here were derived from AlphaFold2-predicted structures rather than experimental structures.

The fold-concordance assessment showed that four domain instances (three Fibronectin type III repeats and one Laminin EGF-like domain) fell below the fold-recovery threshold, and the transmembrane and uncharacterised regions could not explicitly be assessed for lack of experimental templates. Structural features for variants in these regions therefore carry greater uncertainty, and residual inaccuracies in predicted side-chain conformations or loop geometries may introduce additional noise even within well-recovered folds^22,68^.

Although features capturing predicted protein-protein interaction sites (CSM-potential^44^ and ANCHOR2^43^) were incorporated during feature generation, they did not survive the sequential feature selection pipeline, meaning there’s a general lack of features that explicitly account for interaction interface geometry. Variants at these interfaces may be pathogenic through mechanisms orthogonal to local conservation and stability, contributing to the residual false negative rate observed here. Thus, future work should seek to expand the training data with additional functionally characterised *USH2A* variants as they become available, and to incorporate features capturing inter-domain contacts and homomerisation interfaces, informed by emerging structural and proteomic data^65^. Finally, integrating UshEffect-3D predictions into clinical variant curation workflows, alongside existing genetic and ophthalmic examinations, represents a natural next step towards improving the diagnosis of USH2.

## Conclusion

This study establishes a framework of how gene-specific, biologically informed machine learning can be applied to tackle challenges in variant interpretation. Current state-of-the-art protein structure prediction techniques do not provide ideal solutions for characterising large proteins. By validating the predicted folds with suitable experimental templates, we identified a domain-based strategy that enabled a computationally efficient approach to high-confidence feature generation. A simple, binary classifier trained on these features successfully identified a linear combination of this feature set that effectively differentiates pathogenic and benign variants. However, a deeper look into the model through SHAP and ablation studies, uncovered how in the presence of amino acid sequence-based evolutionary context, structural features in the case of Usherin, do not add further discriminatory signal.

Additional benchmarks against general-purpose variant effect predictors highlights that a gene-specific advantage may not exist in this case. This reinforces the conclusions drawn by the ablation analysis that the dominant evolutionary signal prioritised by our model is also easily captured by predictors trained on proteome-wide datasets. A better calibrated decision threshold favoured by UshEffect-3D nonetheless contributes to more accurate variant classification that we apply to VUS interpretation. We therefore propose an expanded landscape of potential USH2 disease-causing variants to prioritise for clinical review.

## Data Availability

UshEffect-3D predictions for ClinVar VUS along with all possible missense variants within the predicted domains are available in the supplementary file associated with this manuscript (Supplementary_File_S1.csv).

## Ethics Statement

As the study utilized publicly available variant data from ClinVar and LOVD, it was determined to be exempt from **Institutional Review Board (IRB)** approval by the University of Queensland. No patient information was accessed or utilized during this research.

## Acknowledgments

We would like to thank Dr. Ashar Malik and Mrs. Sanjana Tule for their immensely helpful feedback on the methodology.

## Author Contributions

Diya Choudhary: Methodology; formal analysis; writing – original draft; visualization; validation. Stephanie Portelli: Writing – review and editing. David B. Ascher: Conceptualization; Methodology; writing – review and editing;

## Supplementary Information

**Table S1:** Distribution of classified *USH2A* missense variants across Usherin domain types. Variant density is the number of classified variants per 100 residues of that domain type. The pathogenic percentage is the proportion of pathogenic variants among those observed in the domain type; odds ratios compare this against the remainder of the protein (Fisher’s exact test, Benjamini-Hochberg corrected across domain types). The PDZ binding motif and transmembrane segment are excluded as they are too small to support inference. \**q <* 0.05.

| Domain type | Residues | Variants | Density | Pathogenic | Benign | OR (95% CI) | Fisher $q$ |
| --- | --- | --- | --- | --- | --- | --- | --- |
| Laminin N-terminal | 247 | 44 | 17.8 | 37 | 7 | 2.94 (1.29–6.72) | 0.013* |
| Laminin G-like | 371 | 52 | 14.0 | 43 | 9 | 2.68 (1.28–5.60) | 0.013* |
| Laminin EGF-like | 535 | 65 | 12.1 | 48 | 17 | 1.54 (0.86–2.75) | 0.210 |
| Uncharacterised | 812 | 87 | 10.7 | 53 | 34 | 0.79 (0.49–1.26) | 0.331 |
| Fibronectin type-III | 3213 | 372 | 11.6 | 226 | 146 | 0.57 (0.40–0.81) | 0.009* |

**Table S2:**
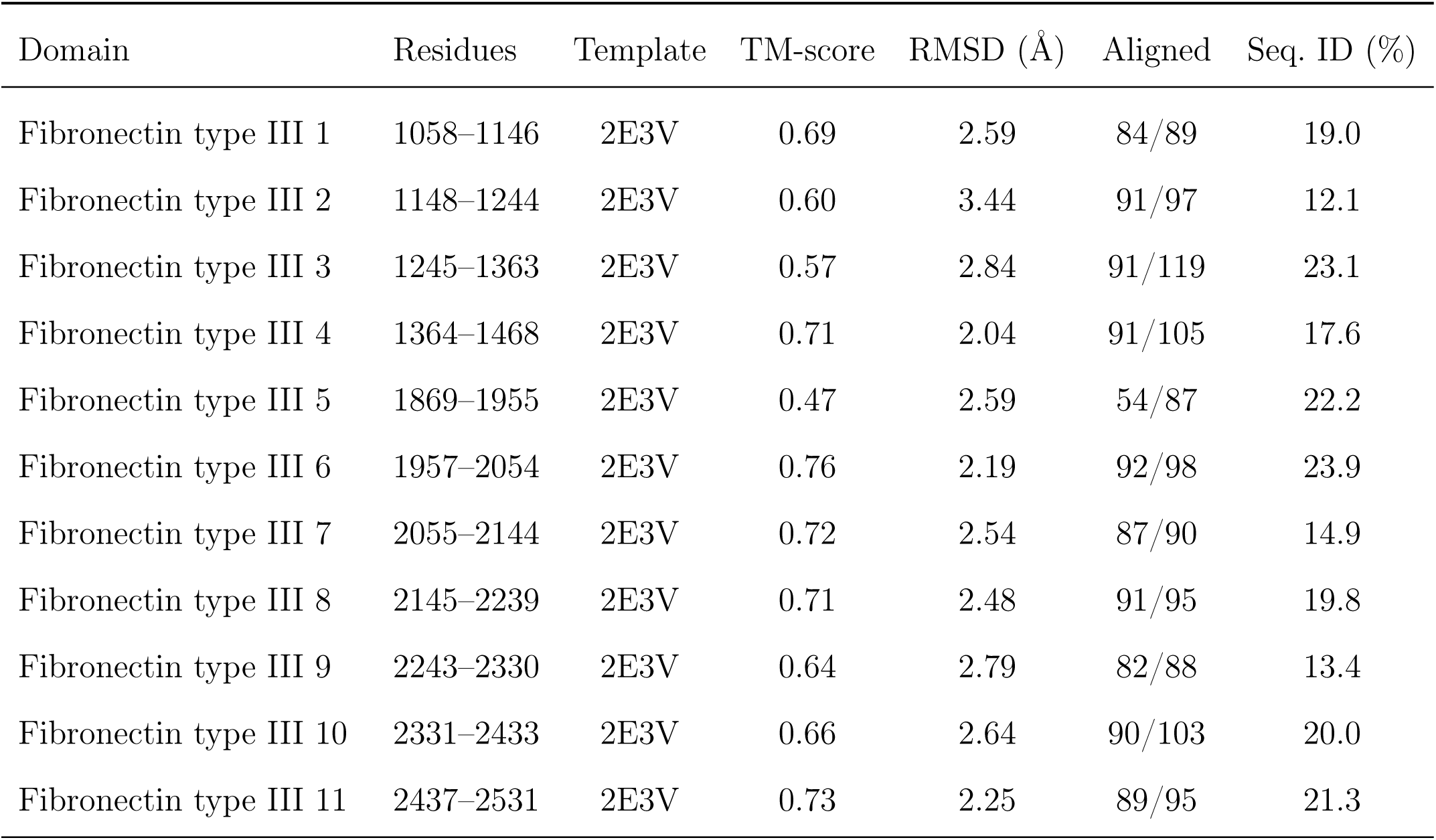
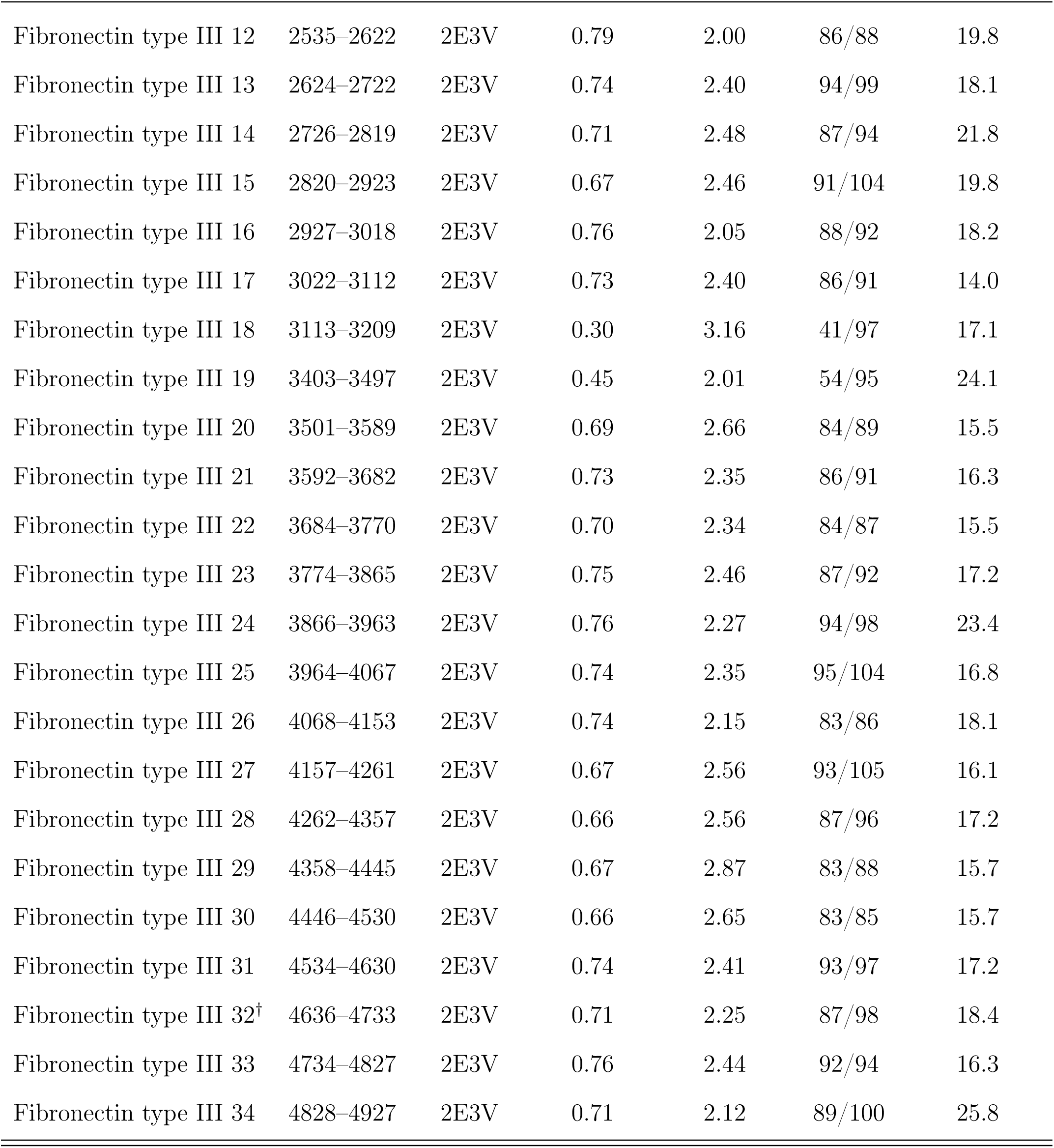
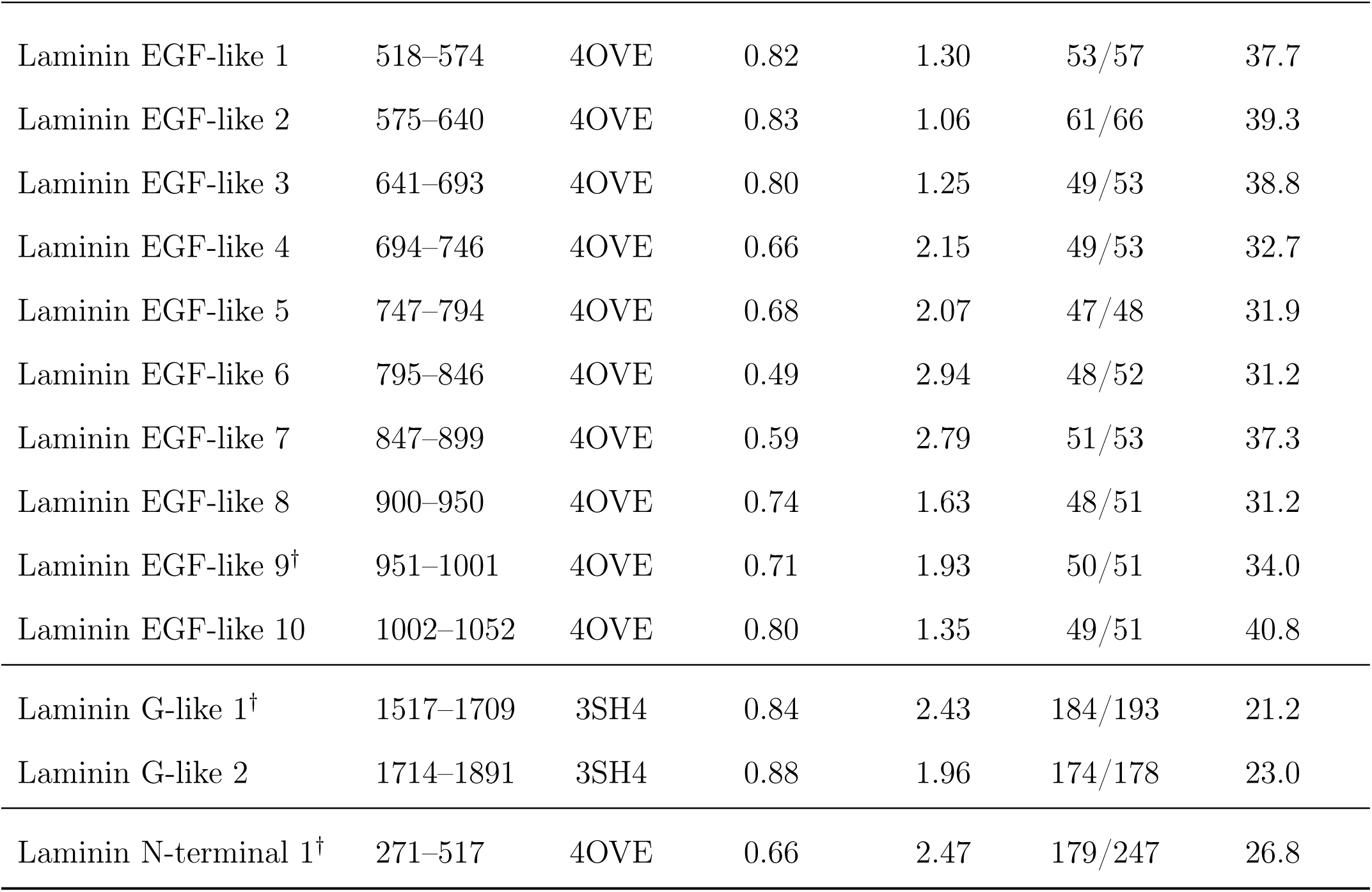
Per-domain fold concordance (TM-align) for all 47 annotated AlphaFold2-derived Usherin domains against a representative experimental structure of each fold family. TM-scores are normalised by predicted-domain length. “Aligned” gives the number of structurally aligned residues over the number of modelled residues in the instance. ^†^Representative instance shown in Figure 2A.

**Table S3:** Nested position-grouped cross-validation performance of all eleven classifiers (mean *±* SD over 10 outer folds). Score (MCC*−*SD) is the model-selection criterion: mean nested-CV MCC minus its fold-to-fold SD, rewarding stability between folds as well as average accuracy. Classifiers are ordered by this criterion, with the selected model listed first.

| Classifier | MCC | Accuracy | BAcc | F1 | Precision | Sensitivity | Specificity | Score (MCC–SD) |
| --- | --- | --- | --- | --- | --- | --- | --- | --- |
| Logistic Regression | **0.77** ± 0.09 | **0.89** ± 0.04 | **0.88** ± 0.05 | **0.91** ± 0.03 | **0.91** ± 0.06 | **0.92** ± 0.05 | 0.84 ± 0.11 | **0.684** |
| SVC | 0.74 ± 0.11 | 0.88 ± 0.05 | 0.87 ± 0.06 | 0.90 ± 0.04 | 0.89 ± 0.06 | 0.91 ± 0.05 | 0.82 ± 0.10 | 0.637 |
| Gaussian NB | 0.74 ± 0.11 | 0.88 ± 0.05 | 0.87 ± 0.06 | 0.90 ± 0.04 | 0.90 ± 0.06 | 0.89 ± 0.05 | **0.85** ± 0.10 | 0.634 |
| AdaBoost | 0.76 ± 0.13 | 0.88 ± 0.06 | 0.87 ± 0.07 | 0.91 ± 0.05 | 0.90 ± 0.07 | 0.92 ± 0.05 | 0.82 ± 0.13 | 0.626 |
| KNN | 0.74 ± 0.11 | 0.87 ± 0.06 | 0.87 ± 0.06 | 0.90 ± 0.05 | 0.90 ± 0.06 | 0.90 ± 0.07 | 0.83 ± 0.10 | 0.626 |
| Extra Trees | 0.74 ± 0.12 | 0.88 ± 0.05 | 0.87 ± 0.06 | 0.90 ± 0.04 | 0.90 ± 0.06 | 0.89 ± 0.04 | 0.85 ± 0.10 | 0.626 |
| XGBoost | 0.76 ± 0.14 | 0.89 ± 0.07 | 0.88 ± 0.08 | 0.91 ± 0.05 | 0.90 ± 0.08 | 0.92 ± 0.04 | 0.84 ± 0.13 | 0.621 |
| MLP | 0.73 ± 0.11 | 0.87 ± 0.05 | 0.86 ± 0.06 | 0.89 ± 0.04 | 0.89 ± 0.06 | 0.90 ± 0.07 | 0.83 ± 0.11 | 0.620 |
| Decision Tree | 0.73 ± 0.11 | 0.87 ± 0.05 | 0.86 ± 0.06 | 0.89 ± 0.04 | 0.90 ± 0.06 | 0.89 ± 0.06 | 0.84 ± 0.11 | 0.619 |
| Gradient Boosting | 0.74 ± 0.13 | 0.88 ± 0.06 | 0.87 ± 0.07 | 0.90 ± 0.05 | 0.89 ± 0.07 | 0.91 ± 0.05 | 0.83 ± 0.12 | 0.612 |
| Random Forest | 0.75 ± 0.14 | 0.88 ± 0.07 | 0.87 ± 0.07 | 0.90 ± 0.05 | 0.90 ± 0.07 | 0.90 ± 0.06 | 0.85 ± 0.11 | 0.608 |

**Table S4:** Held-out test-set performance of all eleven tuned classifiers on the group-aware split, in which no residue position is shared with the training set. The held-out set played no part in model selection.

| Classifier | MCC | Accuracy | BAcc | F1 | Precision | Sensitivity | Specificity |
| --- | --- | --- | --- | --- | --- | --- | --- |
| Logistic Regression | 0.75 | 0.88 | 0.87 | 0.91 | 0.89 | 0.93 | 0.81 |
| SVC | 0.73 | 0.87 | 0.86 | 0.90 | 0.87 | 0.93 | 0.79 |
| Gaussian NB | 0.70 | 0.85 | 0.86 | 0.88 | 0.90 | 0.85 | 0.86 |
| AdaBoost | **0.79** | **0.90** | 0.88 | **0.92** | 0.89 | **0.96** | 0.81 |
| KNN | 0.77 | 0.89 | 0.88 | 0.91 | 0.90 | 0.93 | 0.84 |
| Extra Trees | 0.74 | 0.87 | 0.87 | 0.89 | 0.92 | 0.87 | **0.88** |
| XGBoost | 0.75 | 0.88 | 0.87 | 0.91 | 0.89 | 0.93 | 0.81 |
| MLP | 0.74 | 0.87 | 0.87 | 0.89 | 0.91 | 0.88 | 0.86 |
| Decision Tree | 0.70 | 0.85 | 0.86 | 0.88 | 0.90 | 0.85 | 0.86 |
| Gradient Boosting | 0.73 | 0.87 | 0.87 | 0.90 | 0.90 | 0.90 | 0.84 |
| Random Forest | 0.77 | 0.89 | **0.89** | 0.91 | **0.92** | 0.90 | **0.88** |

### Paired differences against general-purpose predictors

**Table S5:** Paired differences (UshEffect-3D *−* comparator) on the held-out test set, with bootstrap 95% confidence intervals (2000 paired resamples). Both predictors are scored on identical resampled variants, so only their disagreements contribute to the interval. A positive value favours UshEffect-3D; **bold** indicates the 95% confidence interval excludes zero. Intervals are uncorrected for the six comparisons made.

| Comparator | ΔMCC | ΔAUROC | ΔSensitivity | ΔSpecificity | ΔBAcc | ΔPrecision | ΔAccuracy | ΔF1 |
| --- | --- | --- | --- | --- | --- | --- | --- | --- |
| SNAP2 | **+0.49 (+0.25, +0.72)** | **+0.204 (+0.116, +0.301)** | **+0.22 (+0.10, +0.35)** | **+0.26 (+0.03, +0.46)** | **+0.24 (+0.11, +0.36)** | **+0.17 (+0.05, +0.29)** | **+0.24 (+0.13, +0.35)** | **+0.20 (+0.11, +0.30)** |
| CPT-1 | +0.12 (−0.03, +0.27) | −0.001 (−0.025, +0.020) | **+0.30 (+0.19, +0.42)** | **−0.19 (−0.31, −0.08)** | +0.06 (−0.03, +0.14) | **−0.11 (−0.20, −0.04)** | **+0.11 (+0.02, +0.20)** | **+0.13 (+0.05, +0.23)** |
| AlphaMissense | +0.11 (−0.05, +0.26) | +0.007 (−0.030, +0.043) | **+0.25 (+0.15, +0.37)** | **−0.16 (−0.30, −0.04)** | +0.05 (−0.04, +0.13) | **−0.09 (−0.18, −0.01)** | +0.09 (+0.00, +0.18) | **+0.11 (+0.02, +0.20)** |
| VESPAI | +0.10 (−0.06, +0.25) | **+0.046 (+0.006, +0.091)** | **+0.12 (+0.03, +0.22)** | −0.05 (−0.18, +0.08) | +0.04 (−0.04, +0.11) | −0.01 (−0.10, +0.06) | +0.05 (−0.02, +0.13) | +0.05 (−0.01, +0.12) |
| ESM-1b | +0.10 (−0.05, +0.24) | **+0.050 (+0.013, +0.093)** | **+0.12 (+0.03, +0.21)** | −0.05 (−0.16, +0.06) | +0.04 (−0.04, +0.11) | −0.01 (−0.08, +0.05) | +0.05 (−0.02, +0.13) | +0.05 (−0.01, +0.12) |
| PolyPhen-2 | +0.09 (−0.04, +0.23) | +0.046 (−0.001, +0.098) | −0.03 (−0.10, +0.04) | **+0.16 (+0.06, +0.28)** | **+0.07 (+0.00, +0.14)** | **+0.08 (+0.02, +0.14)** | +0.05 (−0.02, +0.11) | +0.03 (−0.02, +0.08) |

### Global SHAP feature importance

#### Sensitivity analysis excluding low-confidence structural environments

To assess whether model performance was dependent on uncertain predicted structural environments, residue-level AlphaFold2 confidence was extracted for all labelled variants and performance was recalculated on the subset of held-out variants with pLDDT *≥* 70, and separately after excluding variants in domain instances that fell below the predefined experimental fold-recovery threshold (Table S8). The trained model was left unchanged throughout.

**Table S6:** Global SHAP feature importance for the selected Logistic Regression classifier. Mean absolute SHAP values were computed with LinearExplainer (independent masker over the training data) across the 435 training variants, and give the feature ordering of Figure 4A. Larger values indicate a greater average influence on individual predictions, irrespective of direction.

| Rank | Feature | Family | Mean \|SHAP\| |
| --- | --- | --- | --- |
| 1 | Demask_entropy | Conservation / sequence | 0.1311 |
| 2 | Demask_log2f_var | Conservation / sequence | 0.1298 |
| 3 | delta_PSIC | Conservation / sequence | 0.1075 |
| 4 | Consurf_score | Conservation / sequence | 0.0977 |
| 5 | DDmut | Structure | 0.0755 |
| 6 | Hydrophobic | Structure | 0.0315 |
| 7 | d_WeakPolar | Structure | 0.0233 |
| 8 | d_VDWClash | Structure | 0.0143 |
| 9 | IUPRED_score | Conservation / sequence | 0.0118 |
| 10 | d_Hydrophobic | Structure | 0.0113 |
| 11 | d_Proximal | Structure | 0.0075 |

Both filters removed few variants, indicating that the labelled set is concentrated in confidently modelled regions: 106 of 110 held-out variants (96%) lie at positions with pLDDT *≥* 70, and 90 of 110 (82%) fall in fold-validated domain instances. Model performance remained comparable, and was marginally improved, under both filters: MCC moved from 0.75 (95% CI 0.62–0.87) on the complete test set to 0.78 (0.65–0.90) at pLDDT *≥* 70 and 0.79 (0.65–0.91) in fold-validated domains, with AUROC essentially unchanged (0.96, 0.97 and 0.97). Sensitivity on the ACMG validation set was likewise stable (0.76, 0.75 and 0.74 respectively). All confidence intervals overlap substantially, so these differences are not individually meaningful; the relevant observation is that no filter degraded performance.

The stricter pLDDT *≥* 90 filter retained only 43 held-out variants and, while sensitivity rose to 0.97, MCC fell to 0.73 with a much wider interval (0.44–0.93), reflecting the small number of benign variants remaining. This subset is too small to support inference and is reported only for completeness.

Model performance therefore remained comparable when evaluation was restricted to confidently modelled positions, indicating that the principal conclusions are robust to structural confidence rather than being an artefact of variants in poorly predicted regions. This does not license greater confidence in individual predictions within low-confidence regions, where the structure-derived descriptors remain less reliable.

**Table S7:** Feature-family model comparison: the tuned Logistic Regression classifier refit on each partition of the selected features, evaluated with position-grouped cross-validation (pooled out-of-fold) and on the held-out test set. Values are point estimates with bootstrap 95% confidence intervals (2000 resamples).

| Model | Evaluation | MCC | AUC | Sensitivity | Specificity |
| --- | --- | --- | --- | --- | --- |
| Conservation / sequence | Grouped CV | 0.73 (0.67–0.80) | 0.92 (0.89–0.95) | 0.91 (0.88–0.95) | 0.81 (0.75–0.87) |
|  | Held-out | 0.75 (0.62–0.86) | 0.97 (0.93–0.99) | 0.91 (0.83–0.97) | 0.84 (0.72–0.93) |
| Structure | Grouped CV | 0.45 (0.37–0.53) | 0.78 (0.73–0.82) | 0.60 (0.54–0.66) | 0.85 (0.80–0.90) |
|  | Held-out | 0.50 (0.35–0.63) | 0.72 (0.62–0.81) | 0.57 (0.44–0.68) | 0.93 (0.84–1.00) |
| Sequence + structure | Grouped CV | 0.77 (0.71–0.83) | 0.92 (0.89–0.95) | 0.93 (0.90–0.96) | 0.84 (0.78–0.89) |
|  | Held-out | 0.75 (0.62–0.87) | 0.96 (0.93–0.99) | 0.93 (0.86–0.98) | 0.81 (0.69–0.92) |
| All minus stability | Grouped CV | 0.74 (0.68–0.80) | 0.92 (0.89–0.95) | 0.91 (0.87–0.94) | 0.82 (0.77–0.88) |
|  | Held-out | 0.71 (0.56–0.84) | 0.96 (0.93–0.99) | 0.91 (0.83–0.97) | 0.79 (0.66–0.91) |
| All minus conservation | Grouped CV | 0.42 (0.33–0.50) | 0.77 (0.73–0.82) | 0.60 (0.54–0.66) | 0.83 (0.77–0.88) |
|  | Held-out | 0.45 (0.29–0.60) | 0.73 (0.62–0.82) | 0.57 (0.44–0.68) | 0.88 (0.78–0.97) |

**Table S8:** Evaluation-only sensitivity analysis of UshEffect-3D to AlphaFold2 structural confidence. The trained model is unchanged and only the evaluation sets are filtered, so nothing is optimised against the test data. pLDDT filters act at the residue level; the fold-validated filter retains variants in domain instances that met the TM-align fold-recovery threshold, excluding the four sub-threshold instances and the transmembrane and uncharacterised regions. Values are point estimates with bootstrap 95% confidence intervals (2000 resamples). The ACMG set contains only pathogenic variants, so specificity, precision, MCC and AUROC are undefined (–).

| Evaluation set | Filter | Retained $n$ | Sensitivity | Specificity | Precision | MCC | AUROC |
| --- | --- | --- | --- | --- | --- | --- | --- |
| Held-out test | All variants | 110 | 0.93 (0.86–0.98) | 0.81 (0.69–0.92) | 0.89 (0.80–0.96) | 0.75 (0.62–0.87) | 0.96 (0.93–0.99) |
| | pLDDT $\geq 70$ | 106 | 0.94 (0.87–0.99) | 0.83 (0.71–0.94) | 0.90 (0.82–0.96) | 0.78 (0.65–0.90) | 0.97 (0.93–0.99) |
| | pLDDT $\geq 90$ | 43 | 0.97 (0.90–1.00) | 0.70 (0.40–1.00) | 0.91 (0.81–1.00) | 0.73 (0.44–0.93) | 0.98 (0.94–1.00) |
|  | Fold-validated domains | 90 | 0.96 (0.91–1.00) | 0.80 (0.65–0.93) | 0.88 (0.79–0.96) | 0.79 (0.65–0.91) | 0.97 (0.94–0.99) |
| ACMG validation | All variants | 78 | 0.76 (0.65–0.85) | – | – | – | – |
| | pLDDT $\geq 70$ | 76 | 0.75 (0.66–0.84) | – | – | – | – |
| | pLDDT $\geq 90$ | 43 | 0.81 (0.67–0.91) | – | – | – | – |
|  | Fold-validated domains | 65 | 0.74 (0.63–0.85) | – | – | – | – |

### Distribution of misclassifications across domain types

Predictions from the trained model were pooled across the group-aware held-out test set and the ACMG validation set, and each misclassified variant was assigned to a domain type by its residue position using the UniProtKB domain annotations^7^. Because the ACMG set contains only pathogenic variants, it contributes false negatives alone, whereas the held-out set contributes errors of both kinds. For each domain type, the error rate was computed as the number of errors divided by the number of variants evaluated in that domain type, with a Wilson 95% confidence interval, and compared with the pooled errors in the remainder of the protein by Fisher’s exact test. The resulting *p* values were corrected across domain types by the Benjamini-Hochberg procedure. The two evaluation cohorts were additionally compared with one another by Fisher’s exact test on their false-negative counts among pathogenic variants. No model was refit at any stage of this analysis.

Pooling the held-out test and ACMG validation sets gives 188 evaluated variants and 32 errors: 8 false positives and 5 false negatives on the test set, and 19 false negatives on the pathogenic-only ACMG set. Error rates by domain type ranged from 15.0% in the Fibronectin type III repeats to 25.0% in the Laminin G-like domains (Table S9), but no domain type differed from the remainder of the protein after correction for multiple testing (all Fisher *q* = 1.00).

These counts are reported for completeness rather than as evidence of domain-specific failure. Outside the Fibronectin type III repeats, which supply 120 of the 188 evaluated variants, every domain type contributes fewer than 20 variants and correspondingly wide Wilson intervals spanning 30–40 percentage points; the apparently elevated rate in the Laminin G-like domains, for instance, rests on 4 errors among 16 variants (95% CI 10.2–49.5%). The evaluated sets are therefore too small to resolve domain-level differences in error rate, and we make no claim about them. The better-supported contrast is between evaluation cohorts rather than between domains: the false-negative rate on pathogenic variants was threefold higher in the ACMG cohort than in the held-out test set (*p* = 0.007), which motivates the feature-level analysis of misclassified variants rather than a positional one.

**Table S9:**
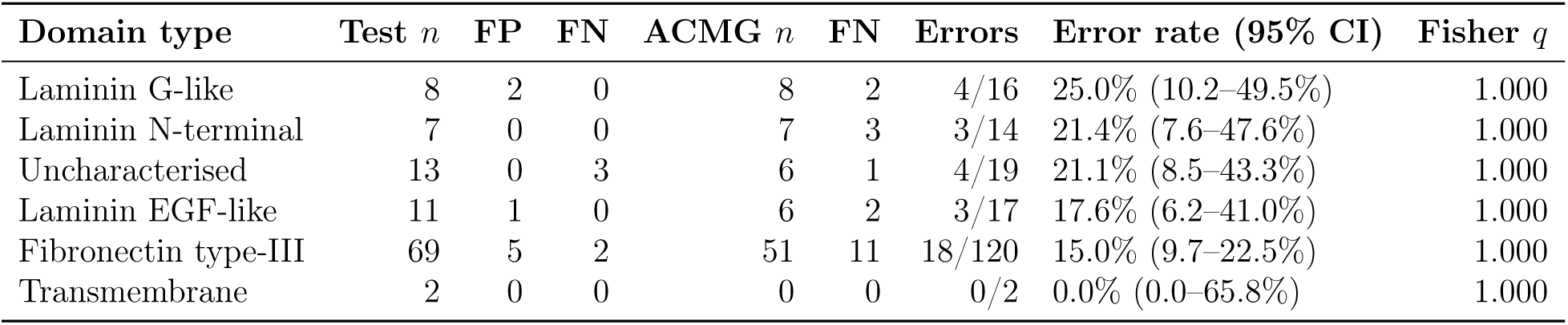
Misclassifications by Usherin domain type across both evaluation sets. The held-out test set contributes false positives and false negatives; the ACMG set is pathogenic-only and contributes false negatives alone. Error rates are pooled across both sets with Wilson 95% confidence intervals, and tested against the remainder of the protein by Fisher’s exact test with Benjamini-Hochberg correction. With only 32 errors in 188 evaluated variants, the intervals are wide and no domain type is distinguishable from any other; these counts are reported for completeness rather than to support a claim of domain-specific failure.

| Domain type | Test $n$ | FP | FN | ACMG $n$ | FN | Errors | Error rate (95% CI) | Fisher $q$ |
| --- | --- | --- | --- | --- | --- | --- | --- | --- |
| Laminin G-like | 8 | 2 | 0 | 8 | 2 | 4/16 | 25.0% (10.2–49.5%) | 1.000 |
| Laminin N-terminal | 7 | 0 | 0 | 7 | 3 | 3/14 | 21.4% (7.6–47.6%) | 1.000 |
| Uncharacterised | 13 | 0 | 3 | 6 | 1 | 4/19 | 21.1% (8.5–43.3%) | 1.000 |
| Laminin EGF-like | 11 | 1 | 0 | 6 | 2 | 3/17 | 17.6% (6.2–41.0%) | 1.000 |
| Fibronectin type-III | 69 | 5 | 2 | 51 | 11 | 18/120 | 15.0% (9.7–22.5%) | 1.000 |
| Transmembrane | 2 | 0 | 0 | 0 | 0 | 0/2 | 0.0% (0.0–65.8%) | 1.000 |

